# Identification and evaluation of small-molecule inhibitors against the dNTPase SAMHD1 *via* a comprehensive screening funnel

**DOI:** 10.1101/2023.01.17.524275

**Authors:** Si Min Zhang, Cynthia B.J. Paulin, Maurice Michel, Petra Marttila, Miriam Yagüe-Capilla, Henri Colyn Bwanika, Huazhang Shu, Rajagopal Papagudi Vekatram, Elisée Wiita, Ann-Sofie Jemth, Ingrid Almlöf, Olga Loseva, Florian Ortis, Christopher Dirks, Tobias Koolmeister, Erika Linde, Sun Lee, Sabin Llona-Minguez, Martin Haraldsson, Kia Strömberg, Evert J. Homan, Martin Scobie, Thomas Lundbäck, Thomas Helleday, Sean G. Rudd

**Author notes:** Corresponding authors: Si Min Zhang & Sean Rudd. C.B.J.P – Research Institutes of Sweden (RISE), Chemical Process & Pharmaceutical Development, Unit Process Chemistry III, 151 36 Södertälje, Sweden. T.L – Mechanistic & Structural Biology, Discovery Sciences, R&D, AstraZeneca, Gothenburg, Sweden.

## Abstract

Sterile alpha motif and histidine-aspartic acid domain containing protein-1 (SAMHD1) is a deoxynucleoside triphosphate (dNTP) triphosphohydrolase central to cellular nucleotide pool homeostasis. Recent literature has also demonstrated how SAMHD1 can detoxify chemotherapy metabolites thereby controlling their clinical responses. To further understand SAMHD1 biology and to investigate the potential of targeting this enzyme as a neoadjuvant to existing chemotherapies we set out to discover selective small molecule-based inhibitors of SAMHD1. Here we report a discovery pipeline encompassing a biochemical screening campaign and a set of complementary biochemical, biophysical, and cell-based readouts for further characterisation of the screen output. The identified hit compound TH6342 and its analogues, accompanied by their inactive negative control analogue TH7126, demonstrated specific, low μM potency in inhibiting the hydrolysis of both natural substrates and nucleotide analogue therapeutics, shown using complementary enzyme-coupled and direct enzymatic activity assays. Their mode of inhibition was subsequently detailed by coupling kinetic studies with thermal shift assays, where TH6342 and analogues were shown to engage with pre-tetrameric SAMHD1 and deter the oligomerisation and allosteric activation of SAMHD1 without occupying nucleotide binding pockets. We further outline the development and application of multiple cellular assays for assessing cellular target engagement and associated functional effects, including CETSA and an in-cell dNTP hydrolase activity assay, which highlighted future optimisation strategies of this chemotype. In summary, with a novel mode of inhibition, TH6342 and analogues broaden the set of tool compounds available in deciphering SAMHD1 enzymology and functions, and furthermore, the discovery pipeline reported herein represents a thorough framework for future SAMHD1 inhibitor development.

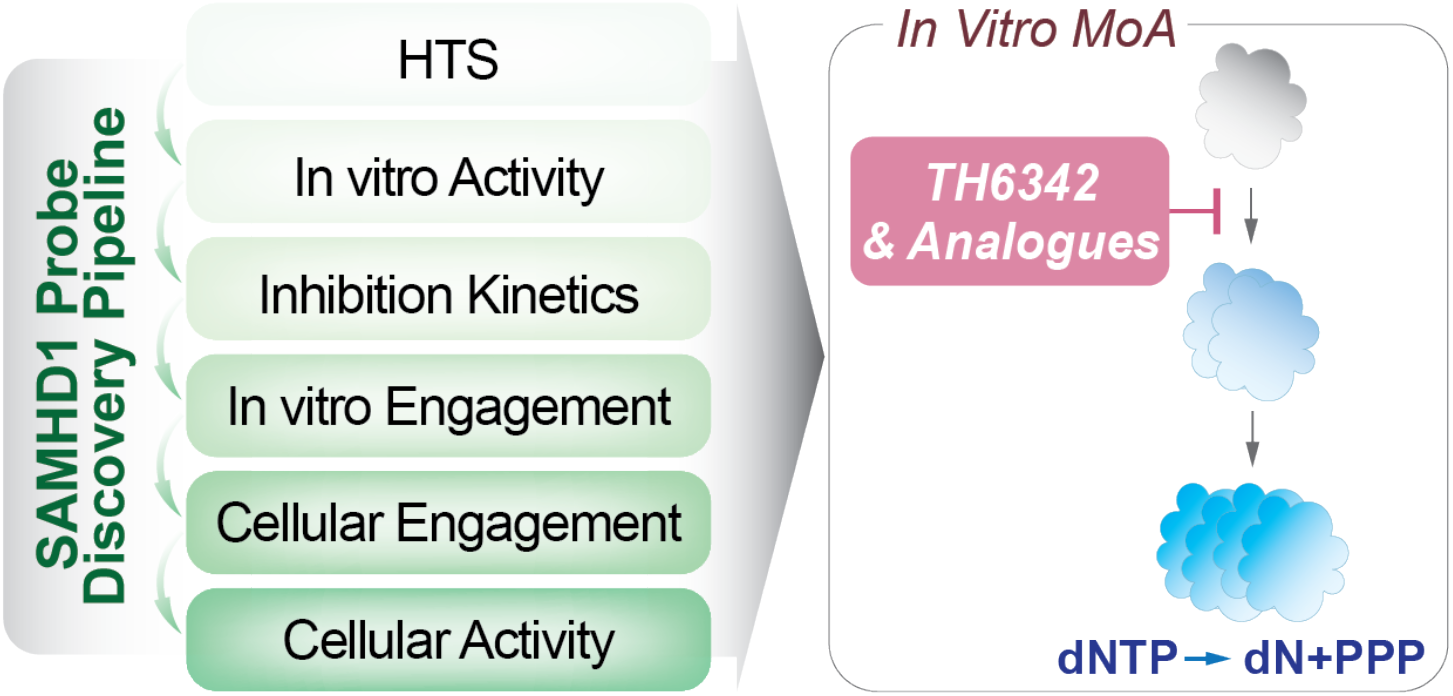

## Introduction

Sterile alpha motif and histidine-aspartic acid domain containing protein-1 (SAMHD1) is a deoxynucleoside triphosphate (dNTP) triphosphohydrolase with critical roles in human health and disease^1,2^. Belonging to the HD-domain superfamily, a group of metal-dependent phosphohydrolases^3^, SAMHD1 catalyses the hydrolysis of canonical dNTPs producing the cognate deoxynucleoside and inorganic triphosphate^4,5^. This activity is regulated allosterically by sequential nucleotide binding, as each SAMHD1 monomer has two allosteric sites with distinct nucleotide binding properties in addition to a catalytic site (reviewed in ref^6^). Allosteric site 1 (AS1) binds to guanine nucleotides such as GTP or dGTP, which promotes formation of the SAMHD1 dimer, whilst allosteric site 2 (AS2) binds any canonical dNTP and promotes formation of the catalytically-competent homotetramer^7-9^. Given that dNTP hydrolysis by SAMHD1 is regulated by the abundance of (d)NTPs, the enzyme acts as a sensor and regulator of cellular nucleotide pool composition^10^.

Adding further to its biological function, SAMHD1 also plays a non-catalytic role in DNA repair, where it is responsible for recruiting enzymes to sites of damage or stalled DNA synthesis^11,12^, and this role is linked to the ability of SAMHD1 to suppress the innate immune response^11,13^. Accordingly, the diverse roles of this enzyme have several implications for human disease. Germ-line mutations in SAMHD1 are associated with the rare hereditary auto-inflammatory disorder Aicardi-Goutières Syndrome^14^ together with early-onset stroke^15^. SAMHD1 mutations are also found in many cancer types, including chronic lymphocytic leukaemia (CLL)^16,17^, T-cell prolymphocytic leukaemia^18^, colon cancer^19^ and mantle cell lymphoma^20-22^ amongst others (recently reviewed in ref^23^).

SAMHD1 was also identified as a Human Immunodeficiency Virus type-1 (HIV-1) restriction factor in myeloid cells^24,25^ and resting T-cells^26^. This is attributed to the ability of SAMHD1 to deplete dNTP pools below the level required for reverse transcription of the viral genome^27-29^, although other functions of SAMHD1, such as nucleic acid binding, have also been shown to contribute^30^. Demonstrating broad antiviral activity, SAMHD1 can also inhibit replication of other retroviruses^31^ and DNA viruses^32^.

In addition to viral restriction, given the dNTP hydrolase activity of SAMHD1 is critical for dNTP pool homeostasis^10^, this has broad implications for cell fitness including maintaining the fidelity of genome duplication^33^ and efficient DNA repair^34^ including class-switch recombination^35^. The dNTP hydrolase activity is also relevant for the activity of nucleobase and nucleoside analogues, a class of chemotherapy that is critical in the treatment of viral infections and cancer^36,37^. These therapies are synthetic mimics of endogenous nucleobases and nucleosides and require bioactivation, typically sequential phosphorylation, inside target cells to exert their anti-viral or anti-cancer properties. The active species of several of these therapies are their triphosphate metabolite and SAMHD1 is capable of hydrolysing a number of these, thus converting them back to their inactive prodrug form^38-44^. Accordingly, SAMHD1 can modulate the efficacy of several of these drugs in disease models^39-49^. In the case of the deoxycytidine analogue cytarabine (ara-C), the standard-of-care in acute myeloid leukaemia (AML), it has been shown to dictate treatment efficacy in the clinic^39,40,50^.

To fully decipher the function(s) of SAMHD1 and their role in various biological processes, and to investigate its therapeutic potential in nucleoside-based oncology treatment^51^, SAMHD1-specific probes with different chemotypes/inhibitory mechanisms are clearly warranted. We and others have previously demonstrated that viral protein-X (Vpx), a simian/human immunodeficiency virus (e.g., SIV and HIV-2) accessory protein, can serve as a biological inhibitor given its ability to target SAMHD1 for proteasomal degradation^39^, albeit challenges remain for its delivery and target specificity^45^. Furthermore, inhibitors of the key nucleotide biosynthetic enzyme ribonucleotide reductase (RNR) can be re-deployed to alter the intracellular nucleotide pool and thereby indirectly suppress the intracellular ara-CTP hydrolysis activity of SAMHD1 in various models of AML^52^, which is now being evaluated in a clinical trial^53^. Previous studies have also explored direct pharmacological inhibition of the dNTP hydrolase activity of SAMHD1^38,41,54-58^, mainly centring around non-hydrolysable nucleoside triphosphates^38,54,57,58^. Whilst the structural similarities to canonical dNTPs allow these molecules to target SAMHD1 in a competitive manner, and thereby broadened our knowledge of SAMHD1 enzymology, their triphosphate moieties prevent good cell permeability and hence limit applications to *in vitro* biochemical studies. Additional past efforts include focused biochemical screening campaigns with a selection of approved therapeutics^55,56^. The identified drugs demonstrated apparent inhibitory activities *in vitro* with IC_50_ values in the 20-100 μM range, but no further studies of their mechanism of action, selectivities or cell activities have been explored^55,56^.

To allow comprehensive SAMHD1 inhibitor characterisation, and to identify potential alternative chemotypes, we established a SAMHD1 inhibitor screening funnel composed of complimentary biochemical, biophysical, and cellular assays, and here report the first tetramer-destabilising non-nucleotide inhibitors. Following the funnel, we first conducted a biochemical screening campaign of a diverse library of 17,656 small molecules, which together with subsequent medicinal chemistry efforts, resulted in a collection of low μM inhibitors (TH6342, TH7127 and TH7528) with specificity for SAMHD1 versus other nucleotide-processing enzymes. We additionally identified a structurally related but inactive analogue TH7126 that represents a suitable negative control. Subsequent mechanistic characterisations via enzymatic and target engagement assays revealed that TH6342 and analogues could deter dimerisation of SAMHD1 and thereby its allosteric activation, representing a novel mode of inhibition. The inhibitor characterisation pipeline was further complemented with in-cell target engagement and SAMHD1 activity reporter assays. Whilst we show that the herein identified chemotype could not inhibit cellular SAMHD1 despite target engagement in cell lysates, TH6342 and analogues, together with competitive SAMHD1 inhibitors as well as viral Vpx protein, constitute a multifaceted set of tools in deciphering SAMHD1 enzymology and functions.

## Results

### Screening campaign for putative SAMHD1 inhibitors

As the first step to develop small-molecule SAMHD1 inhibitors, we screened a small molecule library for potential inhibitors, utilising a previously established and validated enzyme-coupled malachite green (MG) assay (**Fig. 1a**)^39,59^. SAMHD1 produces inorganic triphosphate from dNTP hydrolysis, and this reaction was coupled to that of a pyrophosphatase, which converts the inorganic triphosphate into individual inorganic monophosphates. These can be readily measured using the MG assay, which we used to indirectly determine SAMHD1 activity (**Fig. 1a**). In the screening campaign, dGTP was chosen as the substrate due to its ability to fully activate SAMHD1 through occupying both the AS1 and AS2 sites (**Supplementary Fig. 1 a-b**). Using this assay we screened a library comprising 17,656 distinct chemical entities at a single concentration of 5 μM (conducted by Chemical Biology Consortium Sweden^60^, see **Supplementary Table 1**). Performance of the screening was deemed excellent with an average Z’ factor of 0.75 (**Supplementary Fig. 1c**). Based on a hit criterion of apparent inhibition over three times the standard deviation from the average inhibition of the screening library, 75 hit compounds were identified, yielding a hit rate of 0.42%.

**Fig 1.**
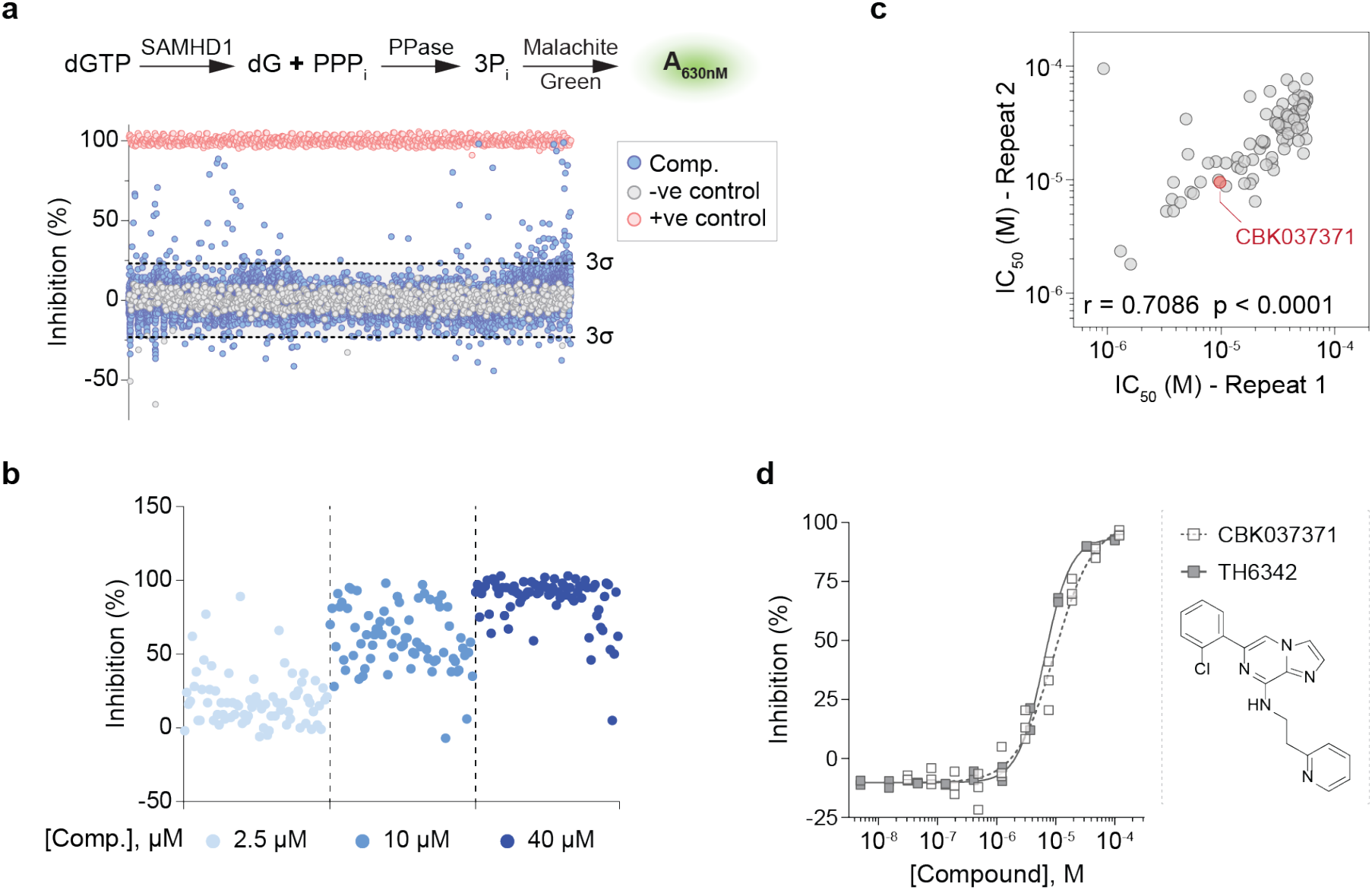
A high-throughput screening campaign to identify putative SAMHD1 inhibitors. **a**. High-throughput screen of 17,656 compounds using the enzyme-coupled malachite green (MG) assay. *Top panel*, schematic presentation of enzyme-coupled MG assay. In the assay, recombinant SAMHD1 was incubated with dGTP and the reaction-released inorganic triphosphate (PPPi) was in turn broken down by inorganic pyrophosphatase (PPase) to inorganic phosphate (Pi) which, following addition of MG reagent, can be quantified by measuring absorbance at 630 nm. *Bottom panel*, scatter plot of SAMHD1 inhibition (%) in the presence of library compounds at 5 μM. Hit identification criteria (dashed line) was defined as three times the standard deviation beyond the average inhibition for the screening library. SAMHD1 only without screening compounds (0% inhibition) and SAMHD1-free (100% inhibition) conditions served as negative and positive controls, respectively. **b**. Confirmation of 75 hit compounds from the screening campaign, exemplified by three-point (2.5, 10 and 40 μM) dose-response inhibitions of recombinant SAMHD1 in the enzyme-coupled MG assay. **c**. Confirmation of 94 selected hit compounds and their analogues *via* two independent 11-point dose-response MG experiments. Concentrations required to inhibit 50% enzymatic activity (IC_50_) from the two experiments agreed, with Spearman correlation r and P values indicated. Compound CBK037371 was selected as the chemistry starting point for further inhibitor development, and is highlighted in red. **d**. In-house synthesised CBK037371 (referred as TH6342 hereinafter) demonstrated similar activity as CBK037371 from screening campaign, shown with 11-point dose-response MG experiments. Individual inhibition % of n = 2-3 independent experiments are shown.

Primary hit compounds were subjected to a round of hit confirmation experiments at multiple concentrations (**Fig. 1b**), and further triaged based on their purity and promiscuity qualities (**Supplementary Fig. 2a**). The resulting 48 final hit compounds, together with a curated library of their in-house available analogues, 200 compounds in total, were then evaluated with an 11-point dose response curve (DRC) test. A subset of 96 compounds with confirmed activity were again re-examined in a second 11-point DRC test. Excellent correlation was observed between the two rounds of DRC confirmation, thus validating the screen output (**Fig. 1c, Supplementary Fig. 2b-c**). CBK037371, a hit compound exhibiting μM inhibitory potency (half maximal inhibitory concentrations, IC_50_ = 9.6 μM), was further validated through in-house re-synthesis and purification (referred to hereafter as TH6342) and was selected for further characterisation as a starting point for medicinal chemistry optimisation (**Fig. 1d**).

**Fig 2.**
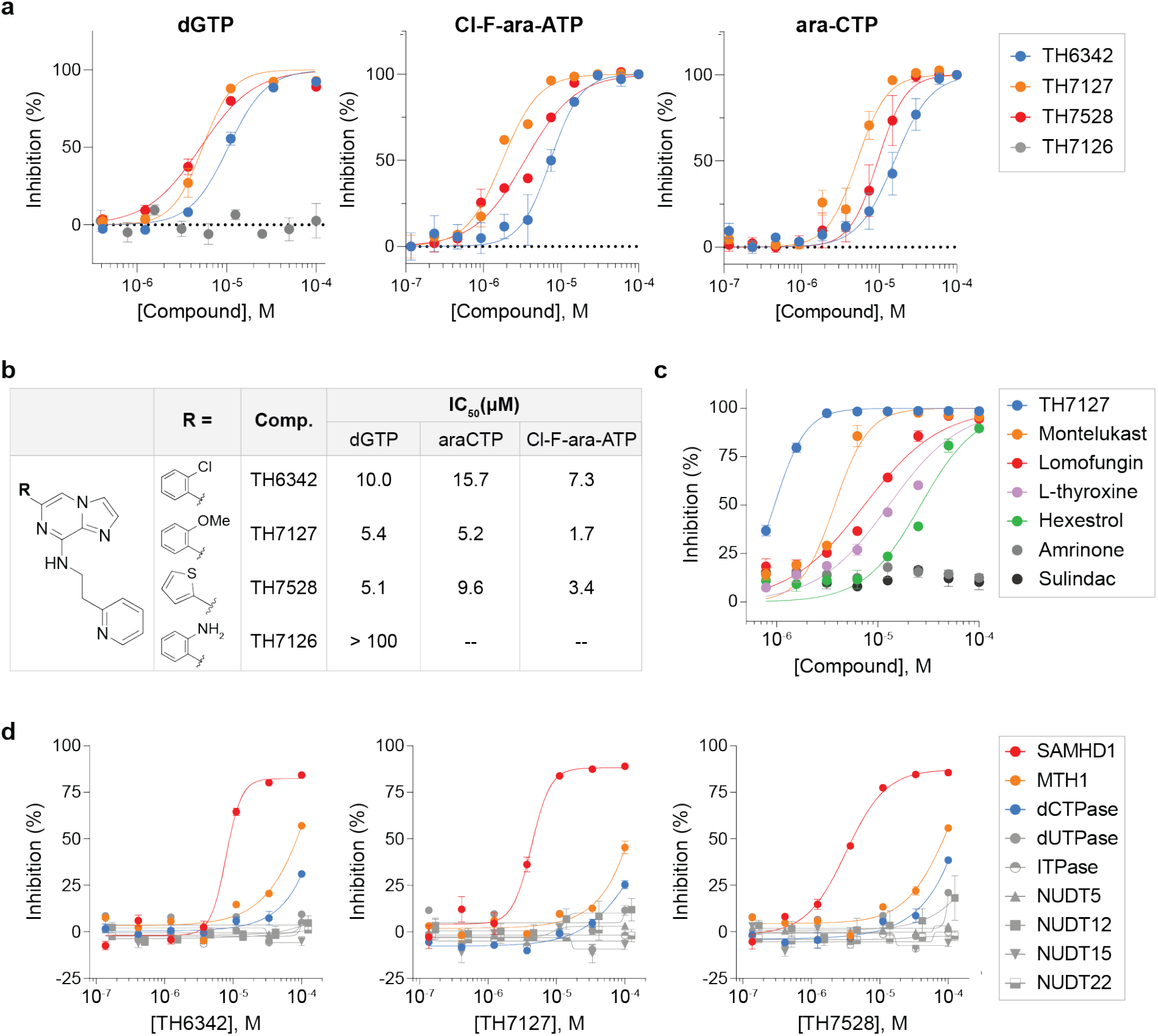
TH6342 and its analogues selectively inhibited SAMHD1 activity with low micromolar potency. **a-b**. TH6342, TH7127 and TH7528, but not TH7126, inhibited the enzymatic activities of SAMHD1 against dGTP, Cl-F-ara-ATP, and ara-CTP. In a, mean inhibition % ± SEM of n = 3 independent experiments are shown. In b, IC_50_ values were determined by curve-fitting mean inhibition % values using a nonlinear regression model (variable slope, GraphPad Prism). **c**. TH7127 demonstrated superior biochemical potency compared to previously published SAMHD1 inhibitors. Mean inhibition % ± SEM of n = 2 independent experiments are shown. **d**. TH6342, TH7127 and TH7528 maintained reasonable selectivity for SAMHD1, when assayed against a panel of nucleoside pyrophosphatases. Mean inhibition % ± SEM of a representative experiment performed in sextuplicate are shown. Enzymatic activities of SAMHD1 in a-d were determined using enzyme-coupled MG assays.

### TH6342 and analogues selectively inhibited SAMHD1

We next initiated a medicinal chemistry follow-up around the initial hit compound TH6342 (see **Supplementary Information** for chemical synthesis), where key chemical features critical for potency were identified by monitoring IC_50_ activities in the MG assay. The structure-activity relationship (SAR) studies (**Supplementary Fig. 3**) were initiated by altering the pyridyl-ethyl-amino part employing a range of primary and secondary amines. A number of similarly active 1,2-diamino or pyridyl-ethyl-amino compounds were generated and confirmed this necessary modification in TH6342. We then directed our attention towards the 2-Chloro-phenyl substituent. Phenyl analogue and heterocycle synthesis led to the development of two compounds with moderately improved activity, TH7127 (2-MeO) and TH7528 (2-thiophenyl), and an inactive control analogue TH7126 (2-Amino). In an attempt to explore the core of the molecule we further synthesised a diverse set of heterocycles with 1,2- and 1,3-disubstitution. Finally, matched pair analysis was performed to build confidence in the series.

**Fig 3.**
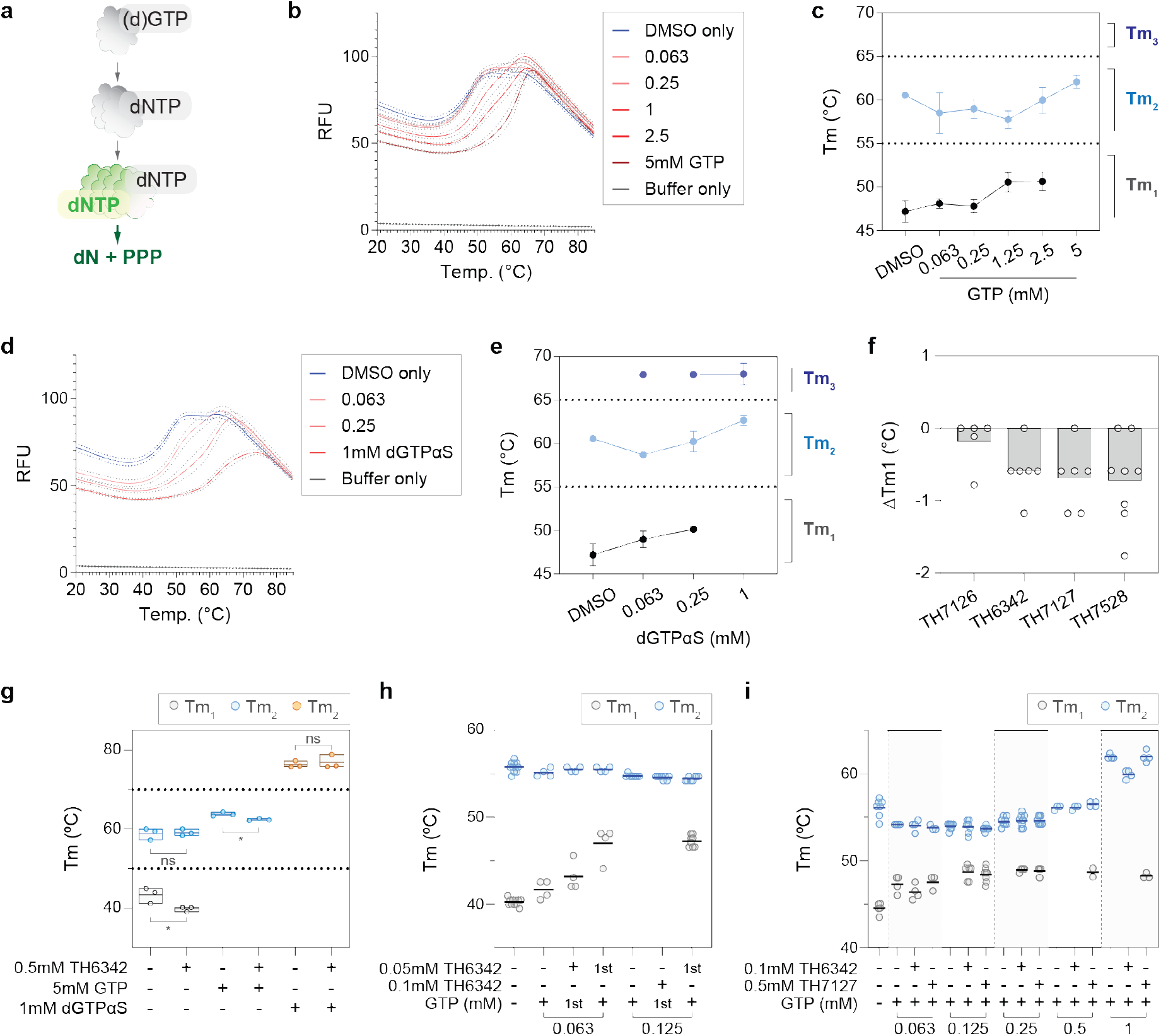
TH6342, TH7127 and TH7528 directly interacted with recombinant SAMHD1 and impeded GTP-induced dimerisation. **a**. Recombinant SAMHD1 protein exhibited two apparent melting temperatures (Tm) in differential scanning fluorometry (DSF) experiments. *Left panel*, schematic representation of ordered activation of SAMHD1. SAMHD1 becomes catalytically competent upon dimerisation as induced by (d)GTP-binding to allosteric site 1 (AS1) and subsequently, tetramerisation by dNTP-binding to allosteric site 2 (AS2). *Right panel*, melting curves (mean relative fluorescence signal ± SEM) of a representative experiment performed in quadruplicate are shown on the left y-axis. Melting temperature were defined by the minima of the negative derivatives of the melting curve (-dRFU/dT), which are shown on the right axis. **b-e**. Melting profile of recombinant SAMHD1 protein in the presence of GTP (b-c) or dGTPαS (d-e). Recombinant SAMHD1 protein was incubated with up to 5 mM GTP (b-c), 1 mM dGTPαS (d-e) or equal volume of DMSO, before its thermal stability being examined by DSF. Mean fluorescence signals (solid line) ± SEM (dashed line) of a representative experiment performed in quadruplicate are shown in b and d. Melting temperatures (Tm) were determined as the minima of negative derivative of the melting curve, and mean Tm ± SD of n = 3-4 independent experiments each performed in quadruplicate are shown in c and e. GTP and dGTPαS delayed the heat-induced denaturation of SAMHD1 and increased the Tm of SAMHD1 in a dose-dependent manner. **f**. TH6342, TH7127 and TH7528, applied at 0.2-0.25 mM, effectively reduced the first apparent melting temperature (Tm_1_) of recombinant SAMHD1 in DSF experiment. Mean changes of Tm_1_ (ΔTm_1_) of n = 2-3 independent experiments performed in triplicates or quadruplicates are shown, where the dots represent individual values of technical repeats in each experiment. **g**. TH6342 at 0.5 mM decreased the Tm of recombinant SAMHD1 protein in the presence of GTP. Mean SAMHD1 melting temperatures of n = 3 independent experiments are shown, together with the individual experiment values. Student’s t tests (unpaired, two-tailed) were performed across treatment groups as indicated in the graph: Tm_1_ (DMSO) Vs. Tm_1_ (TH6342), p = 0.041, t ratio = 2.973, df = 4; Tm_2_ (DMSO) Vs. Tm_2_ (TH6342), p = 0.84, t ratio = 0.2152, df = 4; Tm_2_ (GTP) Vs. Tm_2_ (GTP+TH6342), p = 0.014, t ratio = 4.133, df = 4; Tm_3_ (dGTPαS) Vs. Tm_3_ (dGTPαS+TH6342), p = 0.636, t ratio = 0.5116, df = 4. **h-i**. Incubation with SAMHD1 inhibitors before, but not after GTP treatment deterred SAMHD1 dimerisation. Recombinant SAMHD1 was subject to DSF assay after co-treatment with TH6342 and GTP. In h, SAMHD1 was treated with TH6342 and GTP of comparable concentrations in alternating orders, with compound added first labelled ‘1st’. In i, SAMHD1 was incubated with 0.1 mM TH6342 or 0.5 mM TH7127 prior to increasing concentrations of GTP. Mean SAMHD1 melting temperatures of n = 1-2 (h) or 2 (i) independent experiments are shown, together with the individual experiment values.

Importantly, aside from the natural substrate dGTP, TH6342 as well as its active analogues TH7127 and TH7528 could also inhibit the hydrolysis of ara-CTP and Cl-F-ara-ATP, the active metabolites of the anti-leukemic drugs cytarabine and clofarabine, respectively. Under the same assay conditions, their close analogue TH7126 conferred minimal inhibition, hence serving as a control compound for further mechanistic studies (**Fig. 2a-b**). The low μM potency for TH7127 compared favourably against those of a panel of small-molecule non-nucleotide-based therapeutics previously reported to inhibit SAMHD1 *in vitro*^56^ (**Fig. 2c**). Most critically, we were able to show that TH6342 and the analogues TH7127 and TH7528 selectively inhibited SAMHD1, when assayed up to 100 μM against a panel of nucleotide phosphatases of diverse substrate preferences (**Fig. 2d**). Additionally, the assay systems for these enzymes utilised similar assay conditions such as the choice of coupled enzyme as well as signal detection methodology (**Supplementary Table 2)**, hence offsetting assay interference and further facilitating the selection of SAMHD1-specific inhibitors.

### TH6342 and analogues deterred recombinant SAMHD1 oligomerisation

The binding of TH6342, TH7127 and TH7528 to SAMHD1 *in vitro* was next interrogated using differential scanning fluorimetry (DSF), which evaluates ligand-binding based on target protein thermal stability. To achieve this, we first established assay conditions where we consistently observed stabilisation by known ligands. In the absence of activating-nucleotides, recombinant SAMHD1 mainly displayed two main melting temperatures (Tm) with the first one around 40-45°C (Tm_1_), and the second around 60°C (Tm_2_). SAMHD1 requires sequential nucleotide binding to become catalytically competent, with binding of (d)GTP to AS1 inducing dimerisation and subsequent binding of a dNTP to AS2 to induce tetramer formation (**Fig. 3a**). Addition of GTP to recombinant SAMHD1 (**Fig. 3b-c**) led to a concentration-dependent transition away from the Tm_1_ at 40-45°C to Tm_2_ at 57-60°C, with the latter being the only observed species at 5 mM GTP. Despite variations in the content of folded apo-protein across individual experiments, the observed Tm’s are in good agreement with the known equilibrium between mono- and dimeric species for recombinant SAMHD1^61^, and with recently published thermal shift data for SAMHD1 by Orris *et al*^62^. While the latter study only quotes the first transition between 44-46°C, the figures clearly show two transitions for SAMHD1 in the absence of ligands, which we interpret as representing monomeric (Tm_1_) and dimeric (Tm_2_) species, respectively. Importantly, while the observation of two transitions was not always observed for the apo-protein, signalling a less stable protein in the absence of ligand, this was not the case in presence of nucleotides where we consistently observe ligand-induced stabilisation.

Building on these observations we next performed equivalent experiments replacing GTP with the non-hydrolysable analogue dGTPαS, a known inducer of SAMHD1 tetramerisation. As shown in **Fig. 3d-e**, dGTPαS concentration-dependently stabilised SAMHD1 beyond the first two transitions and elevated its melting temperature to a third Tm_3_ close to 70°C, which likely represents the tetrameric species. The observation of such significant stabilisation is also in agreement with the study by Orris *et al*^62^, although they used elevated dGTPαS concentrations at 2 mM and observed a higher Tm_3_ of 74-76°C across different SAMHD1 forms. Our data clearly shows how dGTPαS is a more potent stabiliser of SAMHD1, as the first (monomer) transition is largely lost already at 0.25 mM concentration.

Having established the Tm shift assay format for recombinant SAMHD1, we subsequently applied this to examine the interactions with inhibitors. In the absence of nucleotides, the inactive analogue TH7126 did not alter SAMHD1 thermal stability at up to 200 μM concentration, while TH6342, TH7127 or TH7528 gave a small concentration-dependent reduction in Tm_1_ (**Fig. 3f, Supplementary Fig. 4**). These small changes clearly signalled that the inhibitors did not behave as the SAMHD1-stabilising nucleotides, *i*.*e*. with a Tm shift towards the higher order species, raising the question whether these compounds inhibited SAMHD1 through irreversible inactivation and/or other mechanisms, e.g. interfering with SAMHD1 multimer stability or formation. To address this, TH6342 and analogues were next subjected to a series of order-of-addition experiments followed by the DSF assay.

**Fig 4.**
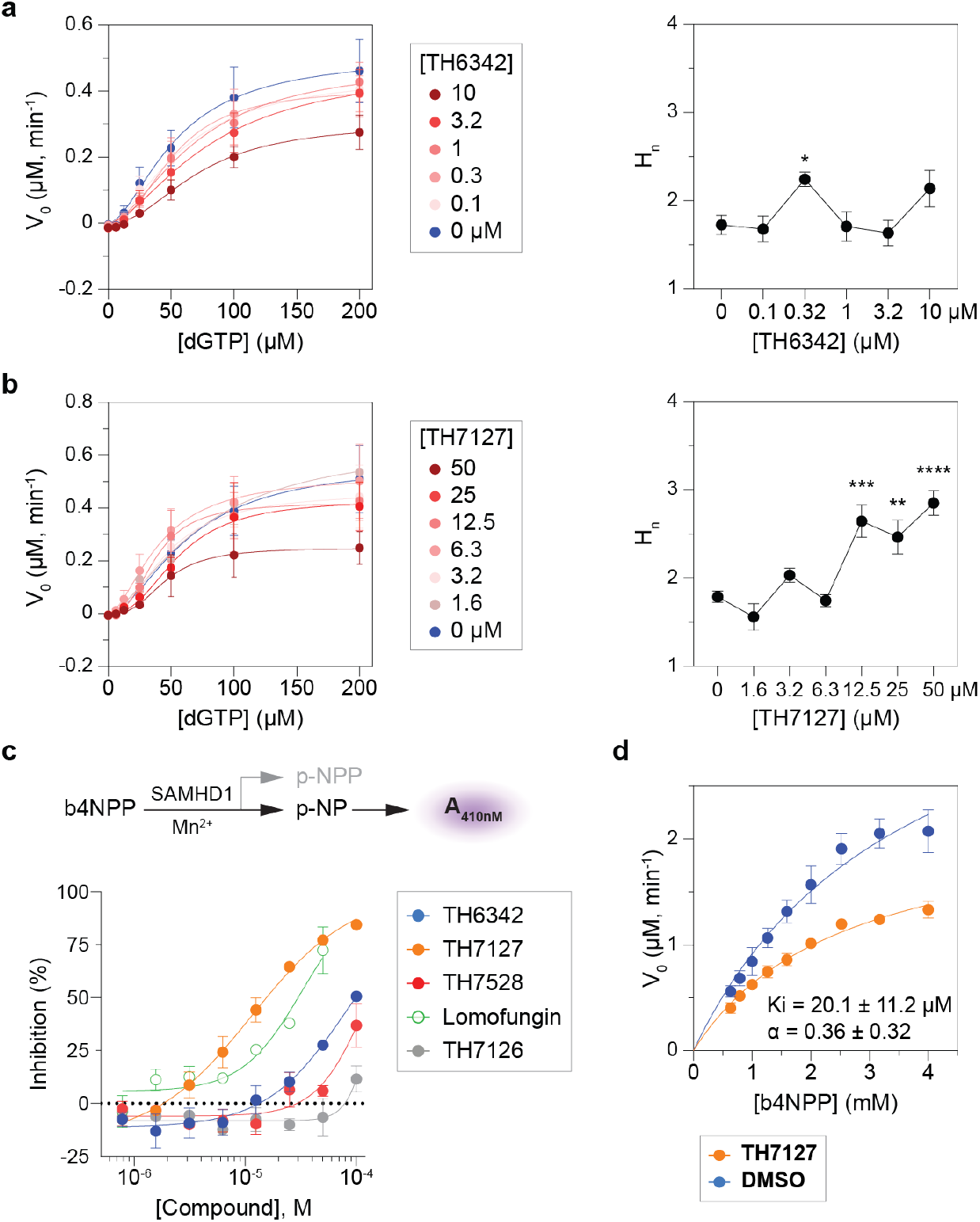
TH6342 and its analogues delayed the allosteric activation of SAMHD1. **a-b**. Study of the inhibition mechanism using the enzyme-coupled MG assay, with dGTP as the substrate. *Left panels*, global fitting of the saturation curves using an allosteric sigmoidal model in GraphPad Prism. *Right panels*, summary of determined Hill coefficient (nH) values. Mean ± SEM of n = 3 (a, b) independent experiments are shown. Fisher’s LSD tests were performed between nH in compound-versus DMSO-treated groups: for TH6342, nH (0 μM) Vs. nH (0.1μM), p = 0.8293, t = 0.2204, DF = 12; nH (0 μM) Vs. nH (0.3 μM), p = 0.0289, t = 2.482, DF=12; nH (0 μM) Vs. nH (1 μM), p = 0.9364, t = 0.08145, DF=12; nH (0 μM) Vs. nH (3.16 μM), p = 0.6672, t = 0.4408, DF=12; nH (0 μM) Vs. nH (10 μM), p = 0.0713, t = 1.979, DF=12. For TH7127, nH (0 μM) Vs. nH (1.56 μM), p = 0.2193, t = 1.275, DF = 17; nH (0 μM) Vs. nH (3.13 μM), p = 0.1588, t = 1.474, DF=17; nH (0 μM) Vs. nH (6.25 μM), p = 0.8094, t = 0.2450, DF=17; nH (0 μM) Vs. nH (12.5 μM), p = 0.0002, t = 4.767, DF=17; nH (0 μM) Vs. nH (25 μM), p = 0.0015, t = 3.765, DF=17; nH (0 μM) Vs. nH (50 μM), p < 0.0001, t = 6.388, DF=17. **c**. TH6342, TH7127, TH7528 inhibited SAMHD1 in a direct SAMHD1 enzymatic assay with b4NPP as the substrate. *Top panel*, schematic representation of the direct SAMHD1 enzymatic assay. In the presence of Mn^2+^ ions, SAMHD1 directly hydrolyses b4NPP into p-NPP and p-NP, with the latter being quantified by absorbance at 410 nm. *Bottom panel*, TH6342, TH7127 and TH7528 inhibited the enzymatic activity of SAMHD1 against b4NPP, with superior activity compared to lomofungin. TH7126 exhibited minimal inhibition of SAMHD1. Mean inhibition % ± SEM of n = 2 independent experiments are shown, and are further fitted with a non-linear regression model (dose-response, variable slope, four parameters, GraphPad Prism). **d**. TH7127 inhibited SAMHD1 following the mixed inhibition model, determined using the direct SAMHD1 enzymatic assay as described in c. Global fitting of the saturation curves, alone or in the presence of TH7127, supported the mixed inhibition model (GraphPad prism). Mean V_0_ ± SEM of n = 2 independent experiments are shown, with Ki and alpha values in the inset.

At high concentrations of stabilising nucleotide, with GTP at 5 mM or dGTPαS at 1 mM, which induce the formation of SAMHD1 dimers and tetramers^61^, subsequent addition of 0.5 mM TH6342 mildly destabilised GTP-, but not dGTPαS-bound SAMHD1 species (**Fig. 3g, Supplementary Fig. 5a**). Considering that SAMHD1 was challenged with a very high concentration of inhibitor compared to its inhibitory potency (∼50x IC_50_), we next co-treated SAMHD1 with reduced levels of GTP and with TH6342 at only 5- and 10-fold above IC_50_, and furthermore at alternating orders. Under these conditions, the incubation with TH6342 following GTP did not significantly affect the protein melting profile (**Fig. 3h, Supplementary Fig. 5b**), signalling the inhibitor does not disrupt already established dimers. On the contrary, when SAMHD1 was first pre-treated with 0.1 mM TH6342 or 0.5 mM TH7127, before the addition of a low concentration of GTP, the allosteric activator was not able to fully remove the Tm_1_ transition, suggesting the persistent presence of monomeric species (**Fig. 3h-i, Supplementary Fig. 5b-d**). For 0.5 mM TH7127 this remained true also when increasing to higher GTP concentrations of 0.5-1 mM, where 0.1 mM TH6342 could not retain the monomeric species (**Fig. 3i, Supplementary Fig. 5c-d**). These data clearly support a model in which TH6342 and analogues inhibited SAMHD1 by hampering GTP-induced dimerisation and thereafter activation, though further kinetic studies were needed to decipher the mechanism of action.

**Fig 5.**
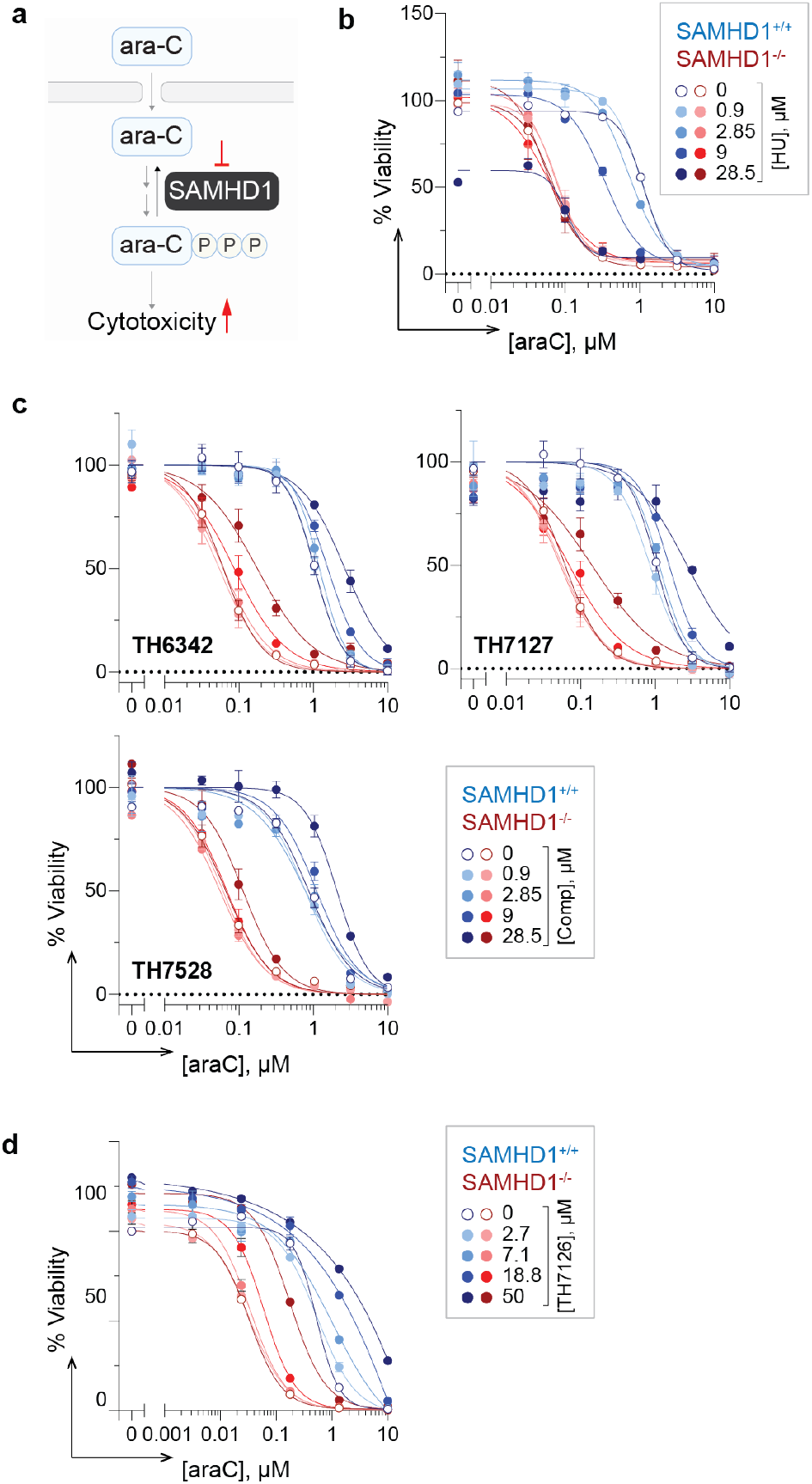
Indirect phenotypic readout to interrogate cellular SAMHD1 inhibition. **a**. Schematic representation of the phenotypic assay to indirectly assess cellular SAMHD1 inhibition. SAMHD1 hydrolyses triphosphorylated ara-C in cells and thereby limits its cytotoxicity, which is re-purposed here to indirectly assay the effects of putative SAMHD1 inhibitors on intracellular SAMHD1. Potential on-target inhibition of cellular SAMHD1 would translate into ara-C potentiation in a SAMHD1-dependent fashion, i.e., only in SAMHD1-competent (SAMHD1^+/+^) but not -deficient (SAMHD1^-/-^) cells. **b-d**. SAMHD1 wildtype (SAMHD1^+/+^) and knockout (SAMHD1^-/-^) THP-1 cells were treated with a concentration matrix of ara-C and hydroxyurea (HU) (b), putative SAMHD1 inhibitors (c), or negative control compound TH7126 (d) for four days, before cell viability was assessed by resazurin reduction assay. Resazurin signals were normalized to DMSO-treated control groups, and mean viability % ± SEM of n ≥ 2 independent experiments each performed in duplicates are shown.

### TH6342 and analogues impeded the allosteric activation of SAMHD1

To delineate the inhibition mechanisms by TH6342 and analogues, we conducted biochemical enzyme kinetic studies using the enzyme-coupled MG assay, with dGTP as the substrate. In agreement with previous studies^54,61^, for reactions applying dGTP as both allosteric activator and substrate, the observed reaction kinetics displayed minimal cooperativity (**Supplementary Fig. 6a**). This can be explained by the relatively rapid allosteric activation to form long-lived SAMHD1 tetramers when compared to the timescale for subsequent steady-state substrate conversion. Interestingly, prior addition of SAMHD1 inhibitors, particularly TH7127, dose-dependently impeded the dGTP-induced SAMHD1 activation, resulting in a shift from a hyperbolic to a sigmoidal dependence on substrate concentration and readily measurable hill coefficients (H_n_) (**Fig. 4a-b, Supplementary Fig. 6b-e**). A similar trend was also observed with TH6342, although less pronounced.

**Fig 6.**
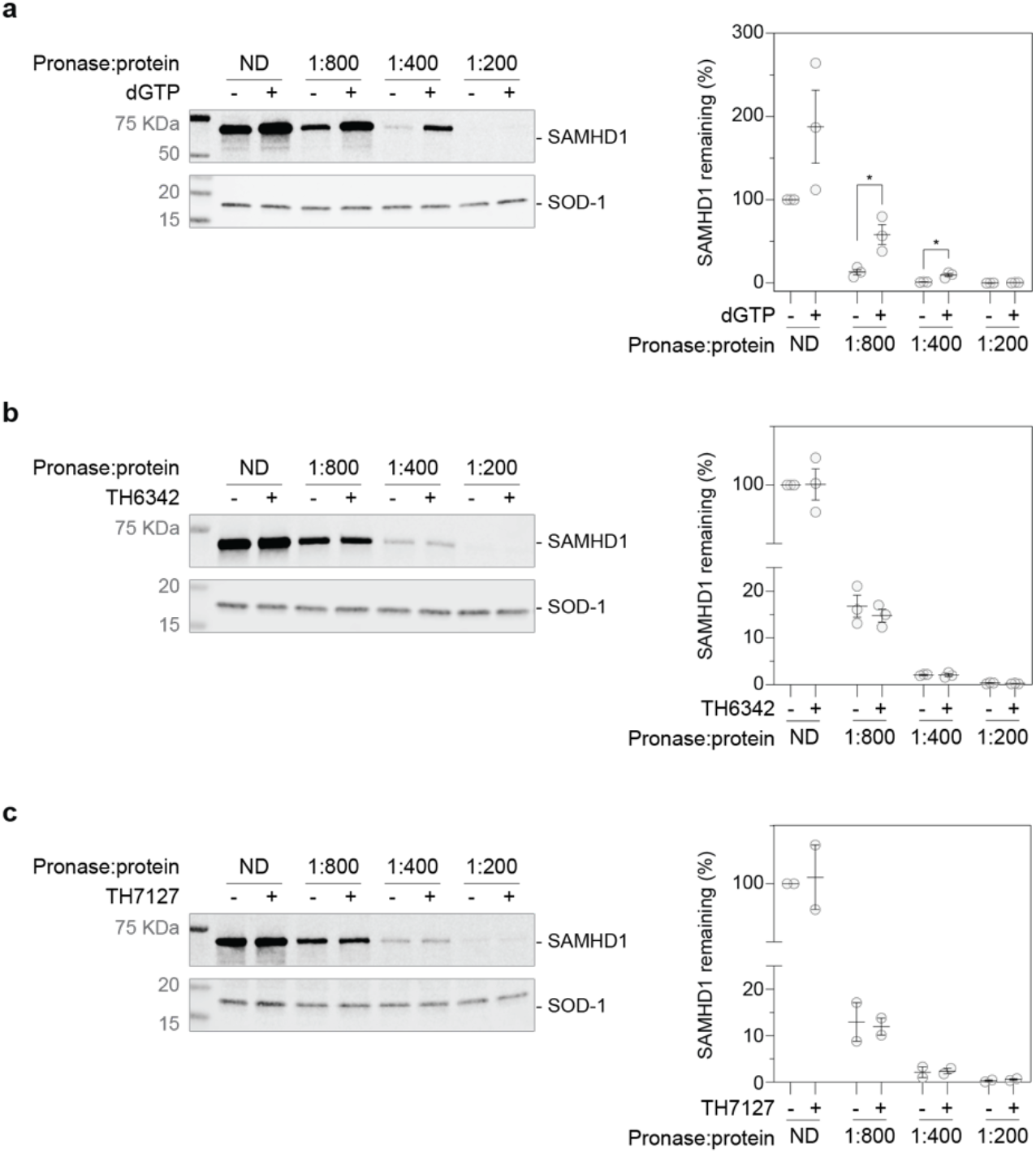
Interrogation of cellular engagement by the putative SAMHD1 inhibitors using the DARTS assay. **a**. Establishment of cellular SAMHD1 engagement DARTS assay using THP-1 cell lysate, with dGTP as the positive control. THP-1 cell lysates were treated with 5 mM dGTP before being digested with pronase at indicated ratios to total protein concentration, and then analysed by Western blot. **b-c**. TH6342 (b) or TH7127 (c) did not show significant engagement of cellular SAMHD1 in THP-1 cell lysate in DARTS assays. THP-1 cell lysate was incubated with 100 μM compound or equivolume of DMSO, before being subject to pronase treatment at indicated enzyme/total protein ratio. Remaining soluble and folded proteins in the cell lysates were then analysed by Western blot. For a-c, *left panels*, representative Western blot images. SOD-1 protein served as the loading control; *right panels*, densitometry analysis, where SAMHD1 signals were normalized to SOD-1 signals and then relative to DMSO control samples received no digestion (ND). Mean relative protein signals ± SEM of n = 3 (a-b) or 2 (c) independent experiments are shown. In a, samples treated with dGTP were compared to DMSO control using student’s t test (unpaired, two-tailed): ND condition, p = 0.117, t = 1.997, df =4; pronase/protein ratio of 1:800, p = 0.022, t = 3.619, df = 4; pronase/protein ratio of 1:400, p = 0.010, t = 4.570, df =4; and pronase/protein ratio of 1:200, p = 0.091, t = 2.219, df = 4.

Data from DSF and enzyme-coupled MG activity assays collectively suggested that the SAMHD1 inhibitors delayed the allosteric oligomerisation of SAMHD1 and thereby its enzymatic activities. To circumvent the influences of allosteric sites and directly address if TH6342 and analogues could also act as competitive inhibitors by binding to the catalytic site, we employed a direct bis(4-nitrophenyl) phosphate (B4NPP) SAMHD1 activity assay. In the assay, sole presence of Mn^2+^ without nucleotides enables SAMHD1 to form catalytically competent active site that can accommodate and hydrolyse B4NPP^56^. Processing of B4NPP results in the formation of yellow p-nitrophenol, thus allowing continuous kinetic measurements (**Fig. 4c-d, Supplementary Fig. 7a-c**). Notably, the alternatively formed active site can also hydrolyse the canonical nucleotide substrates^56^, suggesting no gross structural deviation between the active sites in the direct versus enzyme-coupled MG assays, hence allowing direct interrogation of the effect of TH6342 on the SAMHD1 catalytic pocket. Using this assay, we could confirm that TH6342, TH7127 and TH7528 retained their inhibitory activities (**Fig. 4c**). Notably, TH7127 displayed higher activity than lomofungin, the most potent molecule previously identified using the B4NPP direct enzymatic assay. The inhibition mechanism of TH7127 was subsequently investigated using B4NPP as the substrate. Agreeing with inhibition mechanism suggested by enzyme-coupled MG assay, results from global fitting of the dataset from the B4NPP activity assay did not indicate direct competition for the catalytic pocket, but rather supported a mixed mode of inhibition (Ki = 20.1 μM, alpha = 0.36) (**Fig. 4d, Supplementary Fig. 7d**). Collectively, the data from kinetic studies, as corroborated by the DSF assay, suggested that TH6342 and analogues inhibited SAMHD1 by delaying its allosteric activation.

**Fig 7.**
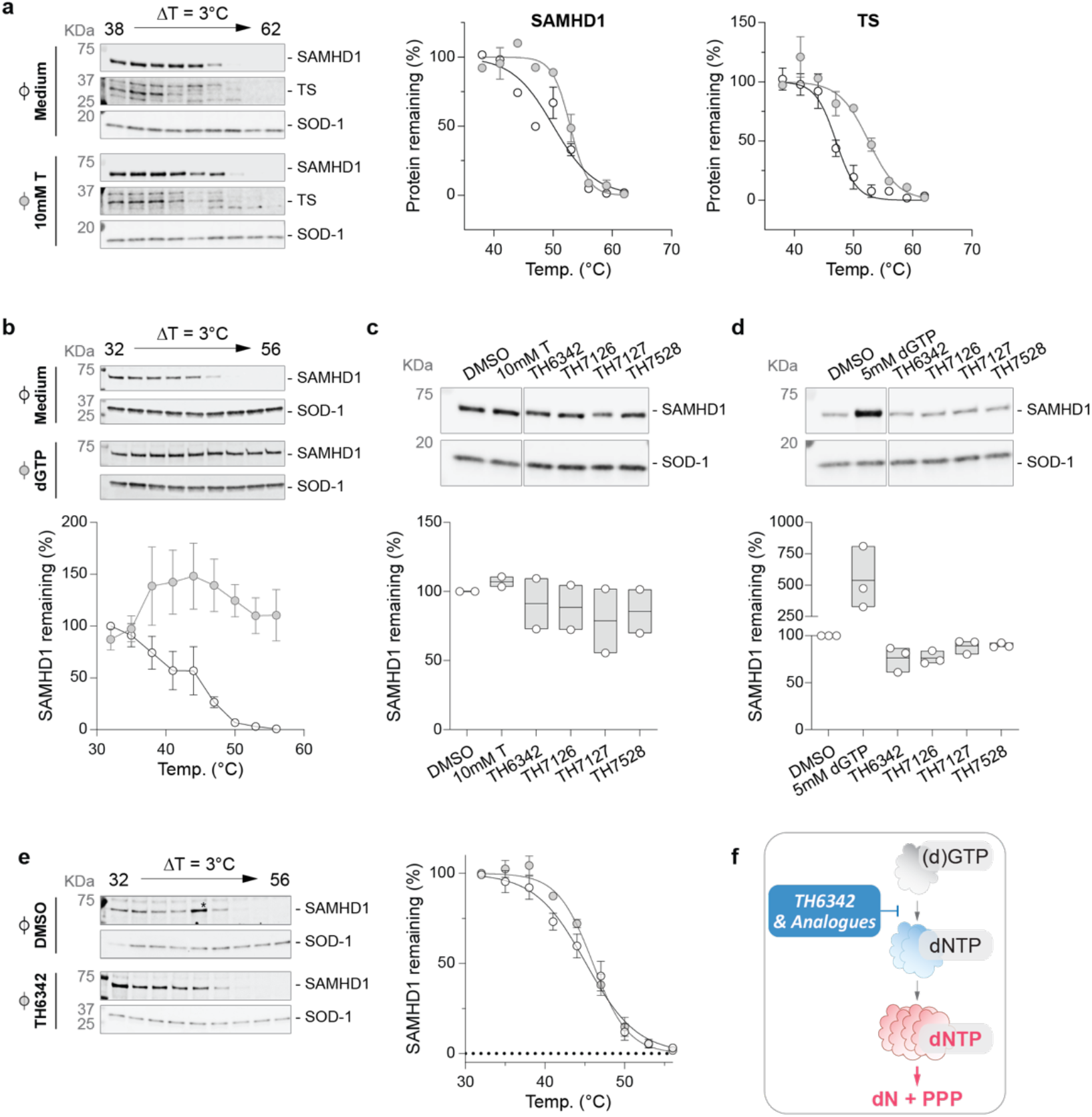
Interrogation of cellular engagement by the putative SAMHD1 inhibitors using CETSA. **a-b**. Establishment of cellular SAMHD1 engagement CETSA assay, validated by thymidine (a) or dGTP (b) as positive controls on intact THP-1 cells (a) or cell lysate (b), respectively. Intact THP-1 cells (a) or cell lysates (b) were treated with 10 mM thymidine (a) or 5 mM dGTP (b), before being heated at indicated temperatures and analysed by Western blot. **c-d**. Engagement of cellular SAMHD1 in intact THP-1 cells (c) or cell lysates (d), interrogated by isothermal single-dose fingerprint CETSA. Intact THP-1 cells (c) or THP-1 cell lysates (d) were treated with 100 μM putative SAMHD1 inhibitors or positive control compounds (10 mM thymidine in c and 5 mM dGTP in d). Following heating at screening temperatures, soluble SAMHD1 was examined *via* Western blot. **e**. TH6342 mildly engaged cellular SAMHD1 in THP-1 cell lysate, shown with CETSA melting curves. THP-1 cell lysate was incubated with 100 μM compound or equivolume of DMSO, before being subject to heating at indicated temperatures. Remaining soluble and folded proteins in the cell lysates were then analysed by Western blot. *Left (a, e) or top (b-d) panels*, representative Western blot images where SOD-1 protein served as the loading control; *right (a, e) or bottom (b-d) panels*, densitometry analysis, where SAMHD1 or thymidine synthase (TS) signals were normalised to SOD-1 signals and then relative to DMSO control samples heated at the lowest temperatures. Mean relative SAMHD1 signal of n = 2 (a-c), 3 (d), or 4 (e) independent experiments are shown with SEM (a, b, e) or values of individual experiments (c-d). **f**. Proposed mechanism of action of TH6342 and analogues. We propose that the chemotypes identified herein, i.e., TH6342 and analogues, directly inhibited SAMHD1 hydrolase activities by deterring the enzyme dimerisation, which is a prerequisite for formation of the catalytically competent SAMHD1 homotetramer.

### Interrogation of cellular activities of the putative SAMHD1 inhibitors

Past *in vitro* studies have identified effective SAMHD1 inhibitors including various nucleotide analogues^38,54,57,58^, still, cellular permeability and target engagement remain challenging, warranting the development of other chemical entities such as small molecule inhibitors. Having demonstrated the *in vitro* potency of TH6342 and analogues, we next interrogated their ability to engage and inhibit cellular SAMHD1.

SAMHD1 has been shown exhaustively to hydrolyse cytotoxic metabolites of nucleoside analogues such as cytarabine^40,45,51,52^. Depletion of SAMHD1, or its indirect inactivation *via* RNR inhibitors (e.g., hydroxyurea), could lead to SAMHD1-dependent sensitisation to these drugs^39,40,52^, which in turn provides opportunities to evaluate cellular activities of putative SAMHD1 inhibitors (**Fig. 5a**). Thus, we utilised a phenotypic assay in which THP-1 acute monocytic leukaemia cells with SAMHD1 knockout or wildtype expression profile^39^, were treated with a dose-response matrix of cytarabine and putative inhibitors for four days before cell viabilities were determined by resazurin reduction assay. Hydroxyurea, which was previously shown to induce SAMHD1-dependent cytarabine sensitisation, served as the control (**Fig. 5b**). Despite their *in vitro* potencies, none of TH6342, TH7127 or TH7528 sensitised THP-1 cells to cytarabine when applied up to around 30 μM, regardless of the SAMHD1 expression status (**Fig. 5c**). In contrast, hydroxyurea dose-dependently sensitised cells to cytarabine in a SAMHD1-dependent manner (**Fig. 5b**). Notably, the inactive control analogue TH7126, when challenged at 50 μM, antagonised cytarabine treatment, suggesting off-target effects at such high concentration tested (**Fig. 5d**).

Inadequate target engagement can often be attributed to poor cellular activities of small molecules, which was not explored in the previous studies of SAMHD1 inhibitors^55,56^. In light of this, we therefore established a series of target engagement assays to examine ligand binding of SAMHD1 in a cellular context, specifically a drug affinity responsive target stability (DARTS) assay and a cellular thermal shift assay (CETSA) (**Fig. 6-7**). DARTS measures on-target binding through monitoring resistance to protease (pronase) digestion, while for CETSA, protein thermal stability. Here, intact cells or cell lysates were treated with potential ligands and then subjected to pronase treatment (DARTS) or heating (CETSA), before remaining folded SAMHD1 was detected *via* Western blot. We could show that known interactors of SAMHD1, thymidine^63^ or dGTP, substantially stabilised SAMHD1 in intact cells or lysates, respectively, against heat- and/or pronase-induced denaturation, thus validating our assay setup (**Fig. 6a, Fig. 7a-b**).

We next screened the SAMHD1-binding potentials of TH6342 and its analogues *via* DARTS assay, where compound-treated cell lysates were subjected to increasing levels of pronase treatment. TH6342 in cell lysates mildly destabilised SAMHD1 to pronase treatment, particularly at lower pronase to protein ratio of 1:800 (**Fig. 6b**), whilst none was observed with TH7127 (**Fig. 6c**). As an orthogonal approach, isothermal single-dose fingerprint CETSA^64^ was next conducted, where compound-treated cells or lysates were subjected to a single screening temperature (**Supplementary Fig. 8a**). Whilst TH6342 applied to the cell lysates mildly destabilised SAMHD1, no engagement was observed in intact cells for any of the tested SAMHD1 inhibitors, indicating low cell permeability (**Fig. 7c-d**). To confirm this, TH6342 was further interrogated in a CETSA melt curve experiment, where SAMHD1 (de)stabilisation was examined over its full melt curve. Data showed that treating cell lysate with 100 μM TH6342 enhanced SAMHD1 thermal stability, albeit mildly (**Fig. 7e**). Altogether, TH6342 demonstrated mild target engagement in cell lysate, still, poor cell permeability and/or insufficient avidity towards SAMHD1 contributed to their low cellular activities, evidenced by the absence of synergy between the inhibitors and cytarabine. The data further highlighted the importance of cellular SAMHD1 engagement assays in characterising intracellular potencies, supporting that it is an integral part of the screening funnel described here for the development of future potent and cell-active SAMHD1 inhibitors.

## Discussion

SAMHD1 has multifaceted roles important to human health and disease, through its enzymatic activity as a central dNTPase and through its non-catalytic activities^2^. More recently, we and others have reported that SAMHD1 can also deactivate antileukemic drugs, notably cytarabine, the backbone therapy for AML, and thereby limit anticancer efficacy^39,40,50^. To thoroughly decipher SAMHD1 biology and investigate its potential as an anticancer target, validated SAMHD1 probes that have undergone systematic and rigorous interrogations are warranted. Previous attempts have focused on screening FDA-approved libraries with limited hit characterisation or dNTP analogue inhibitors that mimic SAMHD1 substrates, however, their dNTP-like moieties disfavour cell permeability and/or high specificity. Here, in this study, we established a multidisciplinary SAMHD1-probe discovery pipeline and identified that a small molecule TH6342, and its analogues, inhibited SAMHD1 hydrolase activity with low-μM potencies and high selectivity, and more interestingly, *via* a novel mode of inhibition that deters efficient allosteric activation of SAMHD1 without occupying (d)NTP binding pockets (**Fig. 7f**).

We initiated this study with one of the largest reported biochemical screening campaigns against SAMHD1, comprised of 17,656 diverse chemical entities of both commercial (Enamine, TimTec, Maybridge and ChemDiv) and/or in-house origins (donation from Biovitrum). SAMHD1 requires subsequent occupancy of its two allosteric sites (AS1 and AS2) by (d)GTP and then any canonical dNTPs, respectively, to dimerise and eventually tetramerise into the catalytically competent species. The formed SAMHD1 tetramers further display impressive long half-life *in vitro*, making neither allosteric site accessible to free ligands^61^, including potential inhibitors. Therefore, in the screening platform, recombinant SAMHD1 protein was incubated with the screening compounds prior to the addition of dGTP, the self-activating substrate, allowing the identification of both competitive inhibitors targeting the catalytic pockets, as well as compounds with alternative inhibitory mechanisms.

We subsequently identified and developed a collection of low-μM, direct, small molecule SAMHD1 inhibitors, TH6342 and its close analogues TH7127 and TH7528, as well as their inactive analogue TH7126 (**Fig. 2c**). TH6342 and analogues further displayed an interesting mode of inhibition. Kinetic study using the MG enzyme-coupled assay with dGTP as the substrate demonstrated that they could dose-dependently increase the Hill coefficient of the reaction, suggesting delayed SAMHD1 activation (**Fig. 4a-b**). We further interrogated the effect of TH6342 and analogues on the catalytic pocket, through an orthogonal B4NPP direct enzymatic assay, where hydrolysis is initiated by Mn^2+^ without the canonical nucleotide-assisted ordered activation. We could confirm that TH6342 and analogues did not target the enzyme catalytic pocket, exemplified by the lack of competitive inhibition in this assay (**Fig. 4d**). Instead, as corroborated by the order-of-addition experiment using DSF assay, pre-treatment of apoenzyme with TH6342 and analogues impeded GTP-induced dimerisation, an essential step of SAMHD1 activation, further suggesting that the chemotypes reported herein delayed SAMHD1 activation by interrupting AS1 occupancy and/or SAMHD1 dimerisation thereafter. Previous studies have developed several series of (deoxy)nucleotide-based SAMHD1 inhibitors for their structural mimicry to the canonical occupants of the allosteric/catalytic sites, such as dNMPNPP and dNTPαS that target the catalytic pocket and the enzyme-substrate complexes^57,58^. Amongst them, the dUTP analogue pppCH2dU inhibits SAMHD1 by delaying its activation, similar to the proposed mechanism of action for TH6342 and analogues (**Fig. 7f**). Yet still, pppCH2dU demonstrated an apparent competitive nature of inhibition as it deters dimer-to-tetramer transition by predominately targeting AS2^54^. We therefore envision that TH6342 and analogues, which specifically inhibited SAMHD1-mediated catalysis through deterring its dimerisation, could complement the tetrameric SAMHD1-targeting dNTP analogue inhibitors, not only as tool compounds to decipher SAMHD1 enzymology, but also to potentially uncover new biological functions of SAMHD1 at different oligomeric states, e.g., as a nucleic acid-binding protein in its monomeric state^65^.

Previous studies of SAMHD1 inhibitors exclusively focus on their utilities in *in vitro* studies with recombinant protein, inadequately addressing their behaviours in engaging and inhibiting cellular SAMHD1, partly due to the lack of appropriate assay systems. In this study, we further embarked on establishing a series of cellular target engagement assays and an indirect cellular SAMHD1 activity assay (**Fig. 6-7**). The cellular target engagement assays, i.e., CETSA and DARTS assays, assessing the on-target binding both in cell lysates and in whole cells were positively validated by employing previously reported known binders to SAMHD1. We further took advantage of the well-established roles of SAMHD1 in hydrolysing and thereby deactivating cytotoxic drugs (e.g., ara-CTP, the active metabolite of cytarabine) and re-purposed it as a proxy readout of the intracellular dNTPase activity of SAMHD1. Low SAMHD1 dNTPase activity, as indirectly elicited by hydroxyurea treatment^52^, is reflected by dose-dependent synergy with cytarabine-induced cytotoxicity (**Fig. 5b**), in line with phenotypes of SAMHD1 abrogation *via* knockout^52^ or introduction of HIV-1 protein Vpx^39^. The intracellular assay systems described here could therefore provide a structured way to evaluate intracellular behaviours of SAMHD1 inhibitors both in this work and in future studies.

Being the first SAMHD1 inhibitors examined for intracellular potencies, TH6342 and analogues engaged cellular SAMHD1 in THP1 cell lysates, though minimally when tested using whole cells. In line with this, they did not synergise with cytarabine, suggesting minimal inhibition of intracellular SAMHD1, despite inhibition of SAMHD1-mediated hydrolysis of ara-CTP *in vitro* (**Fig. 2a-b**). Whilst being the most potent direct SAMHD1 inhibitor of non-nucleotide moiety reported to date, the data indicated that TH6342 and analogues may require improvement in inhibitory potency and/or cell permeability. Nevertheless, cLog*P* values (TH6342, cLog*P* = 3.07; TH7127, cLog*P* = 2.31; and TH7528, cLog*P* = 2.24) predict favourable membrane permeability. Furthermore, SAMHD1 expression varies greatly among cells of different tissue lineages/origins^52^ and is controlled by cell cycle progression^10^. To ensure a high SAMHD1 protein level and thereby large screening window for potential binders, future inhibitor evaluation could be expanded to multiple cell lines of controlled cell cycle status.

In summary, starting with one of the largest screening campaigns against SAMHD1, here we identified and characterised a collection of SAMHD1 inhibitors, i.e., TH6342, TH7127 and TH7528, with low-μM potency and high selectivity as underscored by an inactive analogue TH7126. More intriguingly, these molecules displayed a novel allosteric mode of inhibition that deters efficient SAMHD1 activation. Whilst further improvement is needed for their intracellular potencies, we envision that TH6342 and analogues, together with SAMHD1 competitive inhibitors (e.g., dNTP analogue compounds) as well as the SAMHD1 degradation-inducing viral Vpx protein, constitute a multifaceted set of tools in deciphering SAMHD1 enzymology and functions. Furthermore, this study established a comprehensive screening funnel encompassing biochemical, biophysical, and cell-based functional readouts, providing the community a thorough framework for future SAMHD1 inhibitor identification and development.

## Supporting information

Compiled Supplemental Items

## Acknowledgements

We thank Nicholas Valerie for critical reading of the manuscript and Hanna Axelsson for assistance compiling methods for the screening campaign. Part of this work was facilitated by the Protein Science Facility at Karolinska Institutet/SciLifeLab (http://ki.se/psf), and we acknowledge the National Cancer Institute (NCI), Division of Cancer Treatment and Diagnosis (DCTD), and Developmental Therapeutics Program (DTP) (http://dtp.cancer.gov) for providing Lomofungin.

This project was supported by the Swedish Research Council (2018-02114 to S.G.R., 2015-00162 and 2017-06095 to T.H.), Swedish Cancer Society (19-0056-JIA and 20-0879-Pj to S.G.R., 21-1490-Pj to T.H.), the Swedish Children’s Cancer Fund (PR2019-0014 to S.G.R., PR2021-0030 to T.H., TJ2022-0063 to M.Y-C.), Sjöberg Foundation (2020-2022 to T.H.), the Dr Åke Olsson Foundation for Hematological Research (2022-00304 to S.G.R.), Karolinska Instiutet in the form of a Board of Research Faculty Funded Career Position (to S.G.R.), the Felix Mindus contribution to Leukemia Research (2019-02004 and 2020-02573 to S.M.Z., 2021-01143 to M.Y-C.), the Karolinska Institute foundation for virus research (2020-00249 and 2022-00247 to S.M.Z.), Loo and Hans Osterman Foundation for Medical Research (2022-01262 to S.M.Z. and 2022-01253 to M.Y-C.) and Åke Wibergs Foundation (M22-0011 to S.M.Z). This project has also received funding from Karolinska Institute’s KID funding for doctoral students (to C.D.) and the Innovative Medicines Initiative 2 Joint Undertaking (JU) under grant agreement No 875510 (M.M., E.J.H. E.W.) The JU receives support from the European Union’s Horizon 2020 research and innovation programme and EFPIA and Ontario Institute for Cancer Research, Royal Institution for the Advancement of Learning McGill University, Kungliga Tekniska Hoegskolan, Diamond Light Source Limited. This communication reflects the views of the authors and the JU is not liable for any use that may be made of the information contained herein.

## Author contribution

S.M.Z., C.B.J.P., H.C.B., R.P.V., M.Y-C., H.S., M.M., E.W., P.M., A-S.J., I.A., O.L., C.D., S.L., M.H., K.S., T.L. and S.G.R. designed, performed, and/or analysed biological and biochemical experiments; C.B.J.P., M.M., T.K., S.L-M., E.L., F.O., E.J.H. and M.S. designed, performed, and/or analysed medicinal chemistry experiments; S.M.Z. and S.G.R. compiled data and prepared the manuscript which was revised with T.L.; S.M.Z., T.H. and S.G.R supervised the project. All authors discussed results and approved the manuscript.

## Conflict of interest

The authors have no competing financial interests in relation to the work described.

## Methods and Materials

### Recombinant protein production and purification

Recombinant human SAMHD1 was expressed and purified as described before^39^. Briefly, SAMHD1 was expressed from pET28a(+) (Novagen) vectors expressed with an N-terminal His-tag in *E*.*coli* BL21 (DE3) cells. It was then purified first using HisTrap HP (GE Healthcare) followed by SP cation-exchange (GE Healthcare) or HiLoad 16/60 Superdex 200 (GE Healthcare) columns (See **Supplementary Fig. 9** for protein purity). Recombinant human MTH1, dCTPase, dUTPase, ITPase, NUDT5, NUDT12, NUDT15 and NUDT22 were expressed and purified as described previously^66-69^. Briefly, all proteins were expressed with an N-terminal His-tag in *E*.*coli* BL21 (DE3) cells, where MTH1, dCTPase, dUTPase, NUDT22 were expressed from pET28a(+) (Novagen) vectors, and ITPase, NUDT5, NUDT12 and NUDT15 were expressed from pNIC28 vectors. All proteins, except dCTPase and dUTPase, were purified first using HisTrap HP (GE Healthcare) followed by gel filtration using HILoad 16/60 Superdex 75 (GE Healthcare). dCTPase and dUTPase were first purified using HisTrap HP (GE Healthcare), followed by His tag removal by thrombin (dUTPase) or TEV digestion (dCTPase), and were subsequently further purified using MonoQ ion exchange column.

All proteins were stored at -80°C in storage buffer (20 mM HEPES pH 7.5, 300 mM NaCl, 10% glycerol (v/v) and 0.5 mM TCEP) until further analysis. The Protein Science Facility (Karolinska Institutet, Sweden) performed expression and purification of all proteins, except for dCTPase and dUTPase.

### Reagents (cell culture, drugs, and antibodies)

#### Cell culture

THP-1, THP-1 ctrl, and THP-1 g2c2 cells were cultured in Iscove’s modified Dulbecco’s medium (GE Healthcare) supplemented with 10% fetal bovine serum and penicillin/streptomycin (100 U/mL and 100μg/mL, respectively), at 37°C with 5% CO_2_ in a humidified incubator. THP-1 cells were purchased from ATCC, and its CRISPR-CAS derivative THP-1 ctrl (SAMHD1^+/+^) and THP-1 g2c2 (SAMHD1^-/-^) were generated and characterized as described previously^39,52^. All cell lines were regularly monitored and tested negative for the presence of mycoplasma using a commercial biochemical test (MycoAlert, Lonza).

#### Drugs and antibodies

GTP (Cat.# G8877), cytarabine (Cat.# C1768), montelukast (Cat.# SML0101), L-thyroxine (Cat.# T2376), hexestrol (Cat.# H7753), sulindac (Cat.# S8139), hydroxyurea (Cat.# H8627), b4NPP (Cat.# N3002), p-NP(Cat.# 1048), sodium phosphate (Cat. # 342483) and thymidine (Cat.# T1895) were purchase from Sigma-Aldrich. Ara-CTP (Cat.# NU-1170), dGTPαS (Cat.# NU-424) and Cl-F-ara-ATP (Cat.# NU-874) were purchased from Jena Bioscience. dGTP (Cat.# 27-1870-04) was purchased from GE Healthcare. Lomofungin (NSC106995) was obtained from National Cancer Institute/Division of Cancer Treatment and Diagnosis/Developmental Therapeutics Program. Amrinone (Cat.# GP7331) was purchased from Glentham Life Sciences. Compounds in solid form are typically prepared as 10-50 mM stock solutions in DMSO and stored at -20°C till further use, except for hydroxyurea, B4NPP and p-NP that were prepared in water. Anti-SAMHD1 antibody was purchased from Abcam (Mouse, OTI1A1, Cat.# ab128107) and used at 1:1000 dilution, or from Bethyl Laboratories Inc. (Rabbit, Cat.# A303-691A) and used at 1:2000 dilution. Both anti-SAMHD1 antibodies detected protein bands with molecular weights corresponding to the molecular weight of SAMHD1 which were absent in the SAMHD1 knockout THP-1 cells (g2-2)^39^. Anti-SOD-1 (G-11, Cat.# sc-17767) and anti-thymidine synthase (F-7, Cat.# sc-376161) antibodies were purchased from Santa Cruz Biotechnology and was used at 1:1000 dilutions. Donkey anti-mouse IgG IRDye 680RD (Cat# 925–68072), donkey anti-mouse IgG IRDye 800CW (Cat# 926-32212), donkey anti-rabbit IgG IRDye 800CW (Cat# 926-32213), and Donkey anti rabbit IgG IRDye 680RD (Cat# 926-68073) were purchased from Li-Cor.

### Small molecule screening campaign

#### Small molecule library composition

The screening campaign comprised a combination of in-house and commercially available libraries, to a total of 17,656 compounds. The commercial compounds originate from Enamine, whereas the in-house libraries were partly donated by Biovitrum AB (the origin and composition have been described previously^70^). Compounds included in the screening set were selected to represent a diverse selection of a larger set of 65,000 compounds, while keeping a certain depth to allow crude SAR studies. The selection was also biased towards lead-like and drug-like profiles with regards to molecular weight, hydrogen-bond donors/acceptors and logarithm of partition coefficient (log P). For long-term storage, the compounds were kept frozen at −20°C as 10mM solutions in DMSO under low-humidity conditions in REMP 96 Storage Tube Racks in a REMP Small-Size Store.

#### Small-molecule SAMHD1 inhibitor screening campaign

The screen for SAMHD1 inhibitors was conducted at Chemical Biology Consortium Sweden (www.cbcs.se), using the enzyme-coupled malachite green assay. Recombinant human SAMHD1 (0.26 μM) was incubated with 25μM dGTP and *E. coli* PPase (12.5U/mL), alone or in combination with screening compounds, in assay buffer (25mM Tris-acetate at pH8.0, 40mM NaCl, 1 mM MgCl_2_, 0.005% Tween 20 and 0.3mM TCEP) at room temperature (RT) for 20 minutes. Termination of the enzymatic reaction was done with 4 mM EDTA. The hydrolysis reaction was then measured by absorbance at 630nm (read time of 0.1s per well, Victor 3 from Perkin Elmer) after incubating with malachite green reagent for a minimum of 8min under agitation.

To facilitate screening, aliquots of the compound stock solutions (10mM in DMSO) were transferred to Labcyte 384 LDV plates (LP-0200) to enable dispensing using an Echo 550 acoustic liquid handler (LabCyte). Compound solutions were then dispensed at 10nl/well directly into columns 1–22 of 384-well assay plates (Nunc 242757), while columns 23 and 24 were reserved for controls (see below). The plates were sealed with a peelable aluminium seal (Agilent 24210–001) using a PlateLoc thermal microplate sealer (Agilent) and kept at - 20°C until used. The screening assay was conducted in a total assay volume of 20μl per well in 384-well assay plates. The final compound concentration in the screen was 5μM with a final DMSO concentration of 0.05% in all wells. On the day of screening, enzyme solution (10μl/well) containing SAMHD1 and *E. coli* PPase and dGTP solution (10μL/well) were added using a FlexDrop IV (Perkin Elmer) to assay plates already containing 10nL of compound solutions. Following 20 minutes incubation at RT, 20 μL EDTA was added using a multidrop 384 (Thermo).

On each assay plate, column 24 contained dGTP and SAMHD1-free reaction buffer, and served as positive control (100% enzyme inhibition), while column 23 contained dGTP and SAMHD1 in reaction buffer but no compound served as negative control (0% inhibition). Raw absorbance values at 630nm were then normalised to negative and positive controls on each individual plate. Hit-limit was identified by the average plus three standard deviations of the library compound responses. Subsequent three-dose (2.5, 10 and 40μM) and 11-point concentration response curves (ranging from 119 μM to 12.5 nM) was conducted using the same assay conditions.

### Enzyme-coupled malachite green assays

#### Determination of SAMHD1 in vitro activity

Enzyme-coupled malachite green assays were used to determine the *in vitro* enzymatic activity of recombinant SAMHD1, performed as previously described^39,59^. Briefly, 0.35 μM recombinant SAMHD1 was incubated with 25 μM dGTP in the reaction buffer (25 mM Tris-acetate pH 8, 40 mM NaCl, 1 mM MgCl_2_, 0.005% Tween-20, 0.3 mM TCEP) fortified with 12.5 U/mL *E. coli* PPase, in the presence or absence of compound treatment, at RT for 20 min. The reaction was then quenched by 7.9 mM EDTA (final concentration 3.95 mM), followed by addition of malachite green reagent (final concentration - 0.5 mM malachite green, 12.9 mM ammonium molybdate, 0.036% Tween-20) and incubation at RT for 20 min. Subsequently, absorption at 630 nm wavelength (A_630nm_) was acquired using a Hidex Sense microplate reader.

#### Evaluation of compound selectivity against SAMHD1

Compound selectivity against recombinant MTH1, dCTPase, dUTPase, ITPase, NUDT5, NUDT12, NUDT15 and NUDT22 were evaluated using the enzyme-coupled malachite green assay as described before^69^. Briefly, recombinant proteins were incubated with corresponding substrates in the reaction buffer (MTH1, NUDT5, NUDT12, NUDT15, dUTPase, and NUDT22 – 100 mM Tris-acetate pH 8, 40 mM NaCl, 10 mM Mg acetate, 0.005% Tween-20 and 1 mM DTT; ITPase – 100 mM Tris-acetate pH 8, 50 mM Mg acetate, 0.005% Tween-20 and 1 mM DTT; dCTPase – 100 mM Tris-acetate pH 8, 100 mM KCl, 10 mM MgCl_2_, 0.005% Tween-20 and 1mM DTT) fortified with corresponding phosphatases, in the presence or absence of compound treatment, for 20-40 min at RT. Malachite green reagent was then added. Following 15-20 min incubation at RT, A_630nm_ was measured. See **Supplementary Table 2** for specific assay conditions.

#### Kinetic study of TH6342 and analogues

Kinetic study of TH6342 and analogues was performed with the enzyme-coupled malachite green assay as described above, with the following modifications. Recombinant SAMHD1 is co-treated with a dose-response concentration matrix of inhibitors (0.1-50 μM) and dGTP (6.25-200 μM), where DMSO levels were controlled at 1% in all wells across the plate. Firstly, the inhibitors and DMSO were dispensed using a D300e Digital Dispenser (Tecan) onto a 384-well plate. Subsequently, 0.35 μM recombinant SAMHD1 and 12.5 U/mL *E. coli* PPase in the reaction buffer were added and incubated for 10 min at RT, following which, dGTP was dispensed and the plate was further incubated for 20 min at RT. The reaction was then quenched by EDTA and the malachite green reagent was subsequently added. Upon incubation at RT for 20 min, A_630nm_ was read in a Hidex Sense microplate reader. Standard curves of inorganic sodium phosphate were constructed to determine amount of phosphate released, and subsequently used to determine reaction kinetics (GraphPad Prism).

### B4NPP direct SAMHD1 enzymatic assay

B4NPP direct SAMHD1 enzymatic assay was performed as described^56^. Briefly, 0.5 μM recombinant SAMHD1 was incubated with compounds or equivolume of DMSO control in the reaction buffer (50 mM Tris-acetate pH 8, 100 mM NaCl, 5 mM MnCl_2_, 0.5% DMSO, 2% glycerol, 0.5 mM TCEP) at RT for 10 min, following which 2 mM b4NPP was added and the reaction was then monitored *via* A_410nm_ using a Hidex Sense microplate reader. Reaction linearity was established by varying concentrations of SAMHD1 or b4NPP, in the absence of compound treatment. P-NP standard curve was established by measuring A_410nm_ of increasing concentrations of p-NP. Kinetic parameters of b4NPP hydrolysis by SAMHD1, in the absence or presence of compound treatment, were determined by globally fitting of the entire dataset using a mixed mode of inhibition (GraphPad Prism).

### Target engagement assays

#### Differential scanning fluorometry (DSF)

DSF assay on recombinant SAMHD1 was conducted as described previously, with minor modifications^69^. Briefly, 5 μM SAMHD1 in the assay buffer (25 mM Tris-acetate pH 7.5, 40 mM NaCl, 1 mM Mg acetate, 0.5 mM TCEP) fortified with Sypro Orange (5X, Invitrogen), at the final volume of 20 μL/well. The assay mixture was then subject to a 20-85/90°C temperature gradient for 20 min, with the fluorescence intensities (RFU) measured every second using a LightCycler 480 Instrument II (Roche Life Science). To assay the effects of compound and/or nucleotide treatments on SAMHD1 thermal stability, SAMHD1 was incubated with compounds or nucleotide for 10-20 min before data collection. When both nucleotides and compounds are present, SAMHD1 was incubated with nucleotides first for 10 min, and subsequently with compounds or equivolume of DMSO for 15 min before data collection. Melting temperatures (Tm) were identified by the minima of the negative first derivative (-*d*RFU/*d*T) of fluorescence intensity (LightCycler 480 Software).

#### Cellular thermal shift assay (CETSA)

For isothermal single-dose fingerprint CETSA using intact cells, THP-1 cells were treated with 10 mM thymidine, 100 μM compounds, or equivolume of DMSO (1%) at 37°C and 5%CO_2_ in a humidified incubator. Two hours post-treatment, cells were collected, washed twice by PBS, and then resuspended in PBS supplemented with cOmplete Mini EDTA-free protease inhibitor cocktail (Roche, Merck) at 20 μL per 10^6^ cells, before the cells were heated at 53°C for 3 min in a Veriti 96-well Thermal Cycler (ABI). Samples were subsequently rested at RT for 3 min, supplemented with 40 μL sample buffer (50 mM HEPES pH 7.5, 5 mM β-glycerophosphate, 0.1 mM Na_3_VO_4_, 10 mM MgCl_2_, 2 mM TCEP, 1X cOmplete, Mini, EDTA-free protease inhibitor cocktail), and then lysed through three freeze-thaw cycles with alternating ethanol/dry ice and 37°C water bath. Finally, lysates were clarified by centrifugation at 17,000g for 20 min and then analysed by Western blot. For CETSA using cell lysates, THP-1 cells were first lysed in the sample buffer at 60 μL per 10^6^ cells, through three freeze-thaw cycles with alternating ethanol/dry ice and 37°C water bath, followed by clarification by centrifugation. Clarified lysates were then treated with 5 mM dGTP, 100 μM compounds, or equivolume of DMSO (1%) at 37°C for 0.5-1 h, before being heated at increasing temperatures (CETSA) or 45°C (isothermal single-dose fingerprint CETSA) for 3 min. Following heating, samples were rested at RT for 3 min, clarified by centrifugation, and finally analysed by Western blot. Protein band intensities were normalised to that of the thermally stable loading control SOD-1, and were curve-fitted *via* a Boltzmann sigmoidal model (GraphPad Prism) to determine the T_agg_.

#### Drug affinity responsive target stability (DARTS)

DARTS assay on THP-1 cell lysates was adapted from a previously reported method^71^. THP-1 cells in logarithmic growth phase were collected, washed twice by PBS, and then lysed for 10 min at RT in M-PER™ mammalian protein extraction reagent (Cat.# 78501, Thermo Scientific) supplemented with 1X cOmplete, Mini EDTA-free protease inhibitor cocktail and Halt™ phosphatase inhibitor cocktail (Cat.# 78426, Thermo Scientific). Lysates were then clarified by centrifugation at 16000 x g for 10 min at 4°C. Protein concentrations of the clarified lysates were subsequently determined by Pierce™ Coomassie plus (Bradford) assay reagent (Thermo Scientific), and samples were diluted in 1x TN buffer (50 mM Tris-HCl pH 8.0, 50 mM NaCl) to a total protein concentration of 1 mg/mL. Thereafter, cell lysates were treated with compounds or equivolume of DMSO (1%) at RT for 1 h, followed by digestion with desired concentration of pronase (Roche) solution, or 1XTN buffer only for non-digested samples, at RT for 30 min. Digestion was stopped by the addition of 4x Laemmli sample buffer (Bio-Rad) supplemented with 100 mM DTT and heating at 95°C for 5 min. Samples were then analysed by Western blot, where protein band intensities were normalised to that of SOD-1, and further compared to non-digested samples.

### Western blot

Centrifugation-clarified cell lysates were mixed in β-mercaptoethanol-supplemented Laemmli buffer (Bio-Rad) and then heated at 99°C for 5-10 min. Samples were subject to sodium dodecyl sulfate-polyacrylamide gel electrophoresis (SDS-PAGE) using 4–15% Mini-PROTEAN TGX gels, and proteins were subsequently transferred to nitrocellulose membranes using a Trans-Blot Turbo machine (Bio-Rad). Following blocking with Odyssey Blocking Buffer (LI-COR), membranes were first probed with primary antibodies at RT for 1 h or 4°C overnight, and then probed with species-appropriate secondary antibodies at RT for 30 min. Between incubations, membranes were washed trice using TBST (Tris-buffered saline, 0.1% Tween 20). Protein bands were visualised using an Odyssey Fc Imaging System (Li-Cor Biosciences), and subsequently analysed using Image Studio Lite (Ver. 5.2, Li-Cor Biosciences). All uncropped western blot images are provided in the source data.

### Proliferation inhibition assay

Proliferation inhibition assay was conducted as described previously^52^. Briefly, compounds of indicated concentrations were spotted into 384-well plates using a D300e Digital Dispenser (Tecan). When applicable, DMSO levels were normalised across the plate at the maximum level of 1%. Cells were then seeded into the compound-containing plates using a Multidrop Combi Reagent Dispenser (Thermo Fisher Scientific), at 1000 cells/well in 50 μL medium (serum levels were kept at 5% to facilitate accurate dispensing). Cells were then incubated at 37°C and 5% CO_2_ in a humidified incubator for 72-96 h, before being incubated with resazurin reagent (final concentration - 0.01 mg/mL for another 4-6 h). Fluorescence signals were subsequently acquired using 530/590 nm (ex/em) filters on a Hidex Sense Microplate Reader and were used to determine relative cell viabilities by normalising to cell-only (100% viability) and medium-only (0% viability) wells. Relative cell viabilities were further curve-fitted using a nonlinear regression model (variable slope, four-parameter, GraphPad Prism), when applicable.

## Notes

### Competing Interest Statement

The authors have declared no competing interest.

