## Supplementary material for "Identification and evaluation of small-molecule inhibitors against the dNTPase SAMHD1 *via* a comprehensive screening funnel": Compiled Supplemental Items

* Corresponding authors:

### Supplementary Items

**Supplementary figures 1-9**

**Uncropped western blots**

**Supplementary table 1-2**

**Supplementary information on chemical synthesis**

**Supplementary Figure 1. Supplemental information of the high-throughput screening campaign for SAMHD1 inhibitors. a.** SAMHD1 hydrolytic activity in MG enzyme-coupled assay required the presence of SAMHD1, PPase, and dGTP. Mean absorbance at 630 nm ± SD of n = 2 independent experiments each performed in duplicate are shown. **b.** SAMHD1-mediated hydrolysis of dGTP into dG, with reaction products confirmed using high performance liquid chromatography (HPLC). HPLC profiles of a representative experiment are shown. **c.** Plotted values of the positive control, negative control, and z' factor for each plate of the high-throughput screening campaign. Absorbance at 630 nm ($A_{630nm}$) of positive (SAMHD1-free, represents 100% inhibition) and negative (SAMHD1 only without screening compounds, represents 0% inhibition) controls are plotted on the left axis, and z' factor values are plotted on the right axis. Mean $A_{630nm}$ ± SD of 26 positive or negative control wells included in each plate are shown.

Supp. Fig. 1

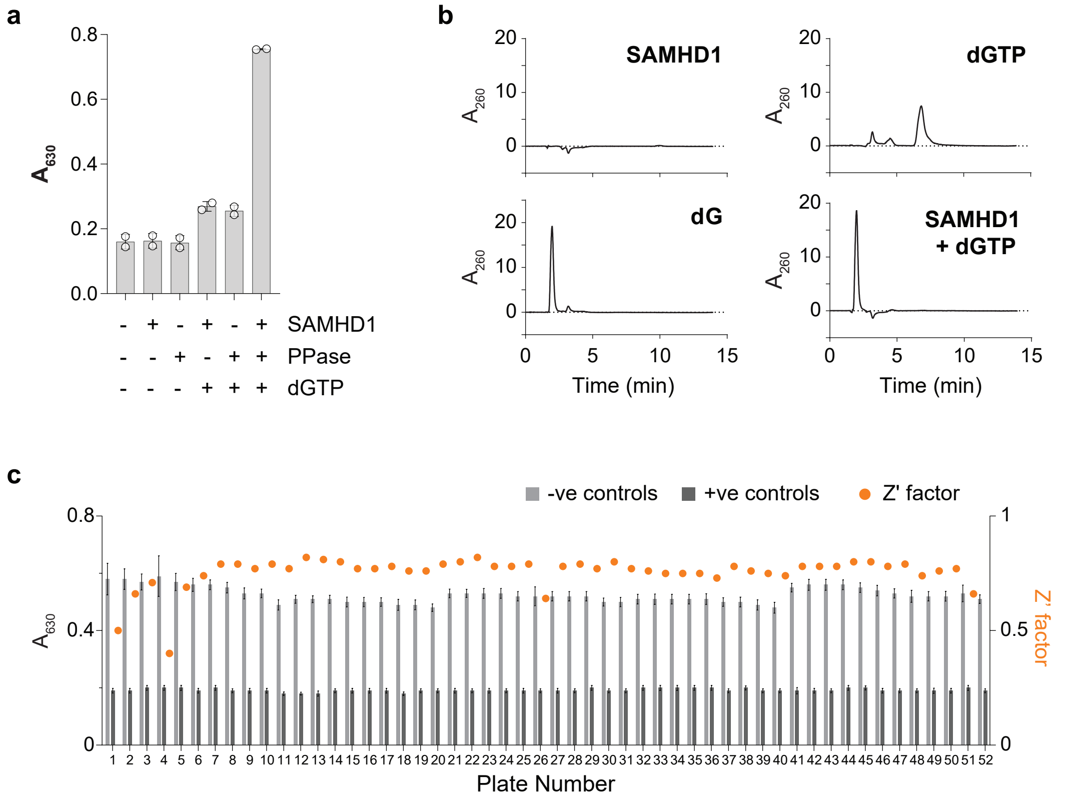

a
0.8
0.6
0.4
0.2
0.0
A630
- + - + - + SAMHD1
- - + - + + PPase
- - - + + + dGTP
b
SAMHD1
dGTP
dG
SAMHD1
+ dGTP
A260
20
10
0
5
15
Time (min)
c
-ve controls
+ve controls
Z' factor
A630
Z' factor
Plate Number

**Supplementary Figure 2. Validation of the high-throughput screening hits. a.** Exclusion of hit compounds displayed promiscuity in previous in-house screening campaigns. For each putative SAMHD1 inhibitor hit, number of times this compound was screened against other targets is plotted on the right axis, and number times this compound was identified as a hit is potted on the left axis. Screening campaigns were summarised based on assay types, i.e., enzymatic, cell-based or engagement assay. **b**. Plotted values of the positive control, negative control, and z' factor for each plate of the 11-point dose-response hit validation. $A_{630nm}$ of positive (SAMHD1-free, represents 100% inhibition) and negative (SAMHD1 only without screening compounds, represents 0% inhibition) controls are plotted on the left axis, and z' factor values are plotted on the right axis. Mean $A_{630nm}$ ± SD of 16 positive or negative control wells included in each plate are shown. **c.** Representative dose-response curve validation of hit compounds. Recombinant SAMHD1 was incubated with increasing concentrations of hit compounds, and enzymatic activities were determined using enzyme-coupled MG assay. Mean inhibition % ± SD of a representative experiment performed in triplicate are shown. The half maximal inhibitory concentrations ($IC_{50}$) were determined by curve-fitting inhibition % using a non-linear regression model (variable slope, four parameters, GraphPad Prisim).

**Supp. Fig. 2**

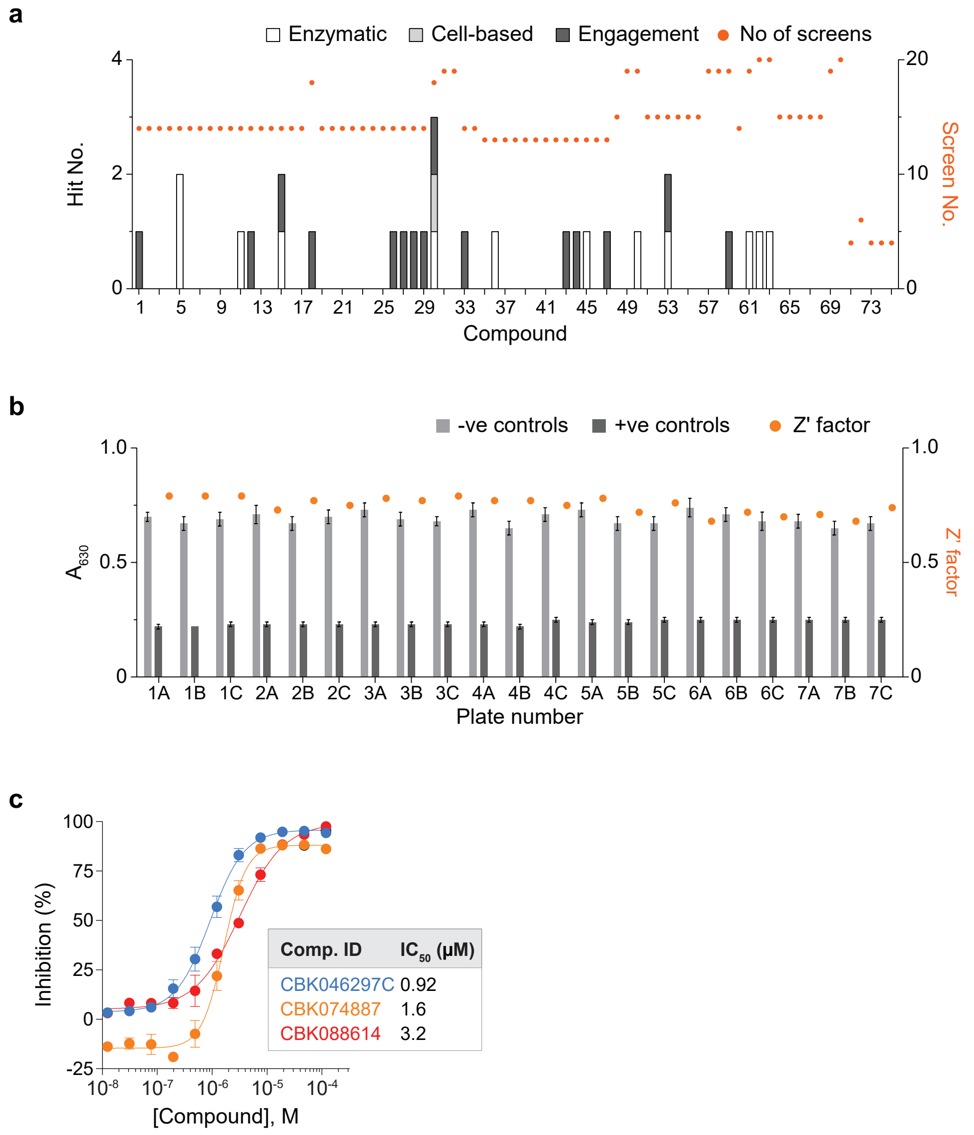
a
Enzymatic
Cell-based
Engagement
No of screens
Hit No.
Screen No.
Compound
b
-ve controls
+ve controls
Z' factor
$A_{630}$
Z' factor
Plate number
1A 1B 1C 2A 2B 2C 3A 3B 3C 4A 4B 4C 5A 5B 5C 6A 6B 6C 7A 7B 7C
c
Inhibition (%)
[Compound], M
Comp. ID | IC$_{50}$ (μM)
CBK046297C 0.92
CBK074887 1.6
CBK088614 3.2

**Supplementary Figure 3. Structure activity relationship (SAR) study of TH6342 and analogues.**

Supp. Fig. 3

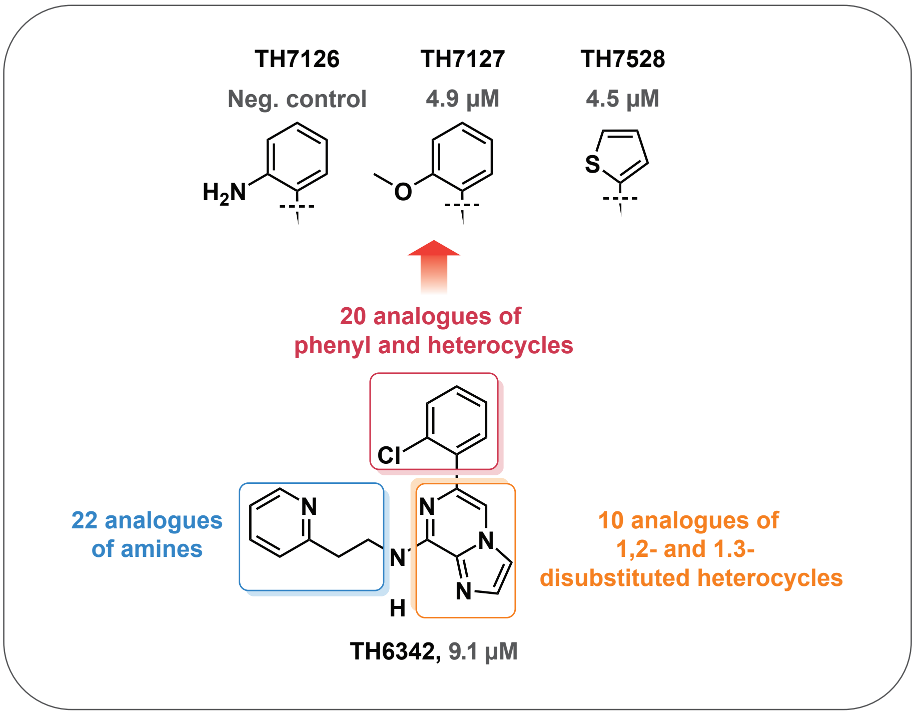

TH7126
Neg. control
TH7127
4.9 µM
TH7528
4.5 µM
20 analogues of
phenyl and heterocycles
22 analogues
of amines
10 analogues of
1,2- and 1.3-
disubstituted heterocycles
TH6342, 9.1 µM

**Supplementary Figure 4. Supplemental information of DSF experiments on recombinant SAMHD1 protein incubated with nucleotide(s) or SAMHD1 inhibitors. a.** Melting curves of recombinant SAMHD1 protein in the presence or absence of putative SAMHD1 inhibitors. Recombinant SAMHD1 protein was incubated with 200-250 μM TH6342, TH7127 and TH7528 or equal volume of DMSO, before its thermal stability being examined by DSF. Mean fluorescence signals (solid line) ± SEM (dashed line) of a representative experiment performed in triplicate are shown. **b-d.** TH6342 (b), TH7127 (c) and TH7528 (d) reduced the $Tm_1$ of recombinant SAMHD1 in a dose-dependent manner. *Left panels*, mean $\Delta Tm_1$ ± SEM of $n \geq 2$ independent experiments performed in triplicates or quadruplicates are shown. *Right panels*, melting curves of recombinant SAMHD1 protein in the presence of TH6342, TH7127, or TH7528. Mean fluorescence signals (solid line) ± SEM (dashed line) of a representative experiment performed in triplicate are shown.

**Supp. Fig. 4**

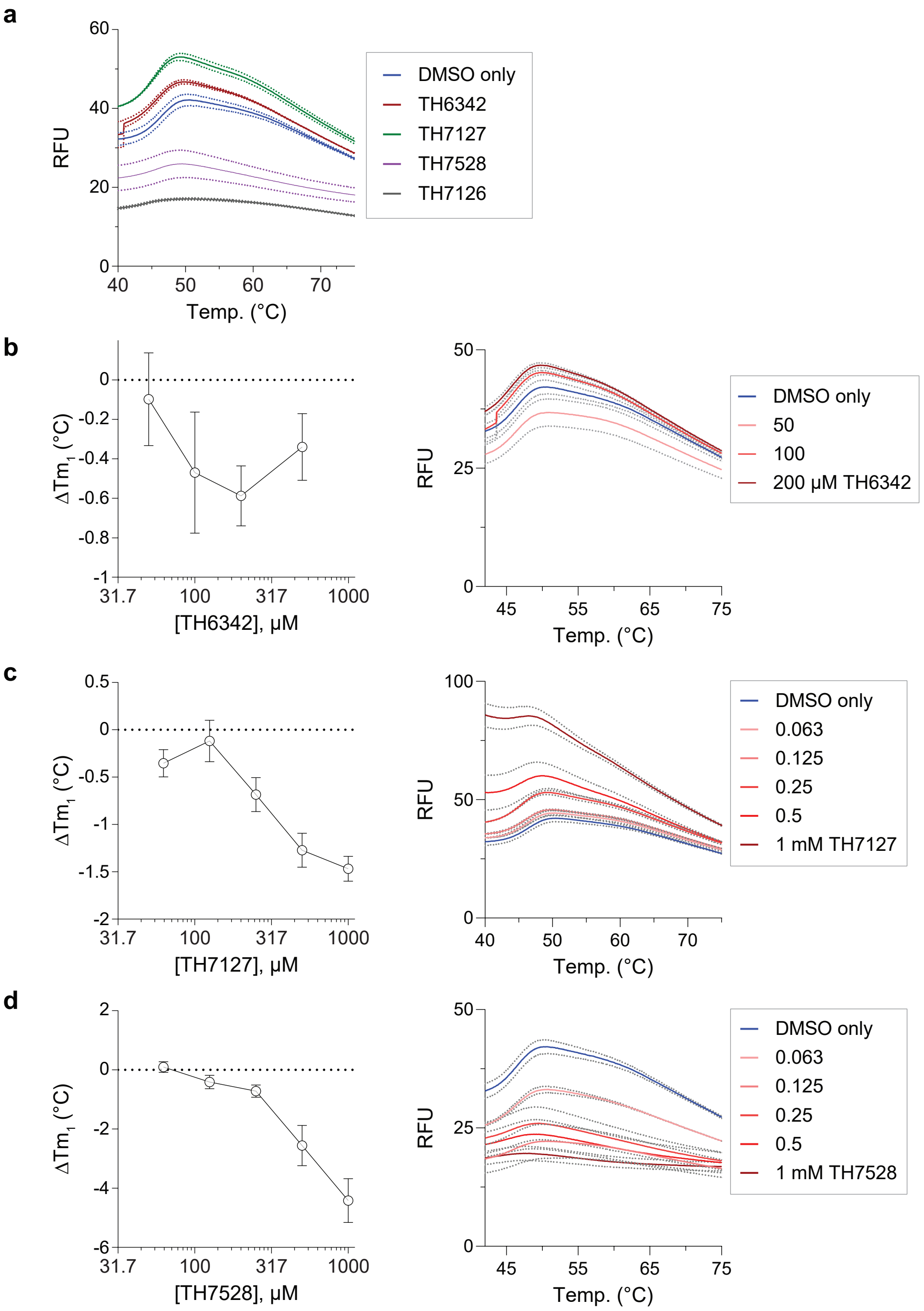
a
RFU
Temp. (°C)
DMSO only
TH6342
TH7127
TH7528
TH7126
b
ΔTm1 (°C)
[TH6342], µM
RFU
Temp. (°C)
DMSO only
50
100
200 µM TH6342
c
ΔTm1 (°C)
[TH7127], µM
RFU
Temp. (°C)
DMSO only
0.063
0.125
0.25
0.5
1 mM TH7127
d
ΔTm1 (°C)
[TH7528], µM
RFU
Temp. (°C)
DMSO only
0.063
0.125
0.25
0.5
1 mM TH7528

**Supplementary Figure 5. Supplemental information of DSF experiments on recombinant SAMHD1 protein co-treated with nucleotide(s) and SAMHD1 inhibitors. a**

TH6342 at 0.5 mM decreased the Tm of recombinant SAMHD1 protein in the presence of GTP. Melting profile of recombinant SAMHD1 treated with activating nucleotides (1 mM GTP or 5 mM dGTPαS), alone or followed by 0.5 mM TH6342. *Left panel*, melting curves of recombinant SAMHD1 in different treatment groups. Mean fluorescence signals (solid line) ± SEM (dashed line) of a representative experiment performed in quadruplicate are shown. *Right panel*, negative derivative (-dRFU/dT) of the SAMHD1 melting curves shown in the left panel are shown. Mean negative derivative values (solid lines) ± SEM (dashed lines) of a representative experiment performed in quadruplicates are shown. **b.** Melting profile of recombinant SAMHD1 co-treated with GTP and TH6342, at alternating orders. *Left panel,* mean fluorescence signals (solid line) ± SEM (dashed line) of n = 2 independent experiments performed in quadruplicate are shown; *right panel,* mean negative derivative (-dRFU/dT) (solid line) ± SEM (dashed line) of the melting curves are shown. **c-d**. Melting profile of recombinant SAMHD1 treated with 0.1mM TH6342 or 0.5 mM TH7127, followed by increasing concentrations of GTP. In c, mean fluorescence signals (solid line) ± SEM (dashed line) of n = 1-2 independent experiments performed in quadruplicate are shown; in d*,* mean negative derivative (-dRFU/dT) (solid line) ± SEM (dashed line) of the melting curves are shown.

**Supp. Fig. 5**

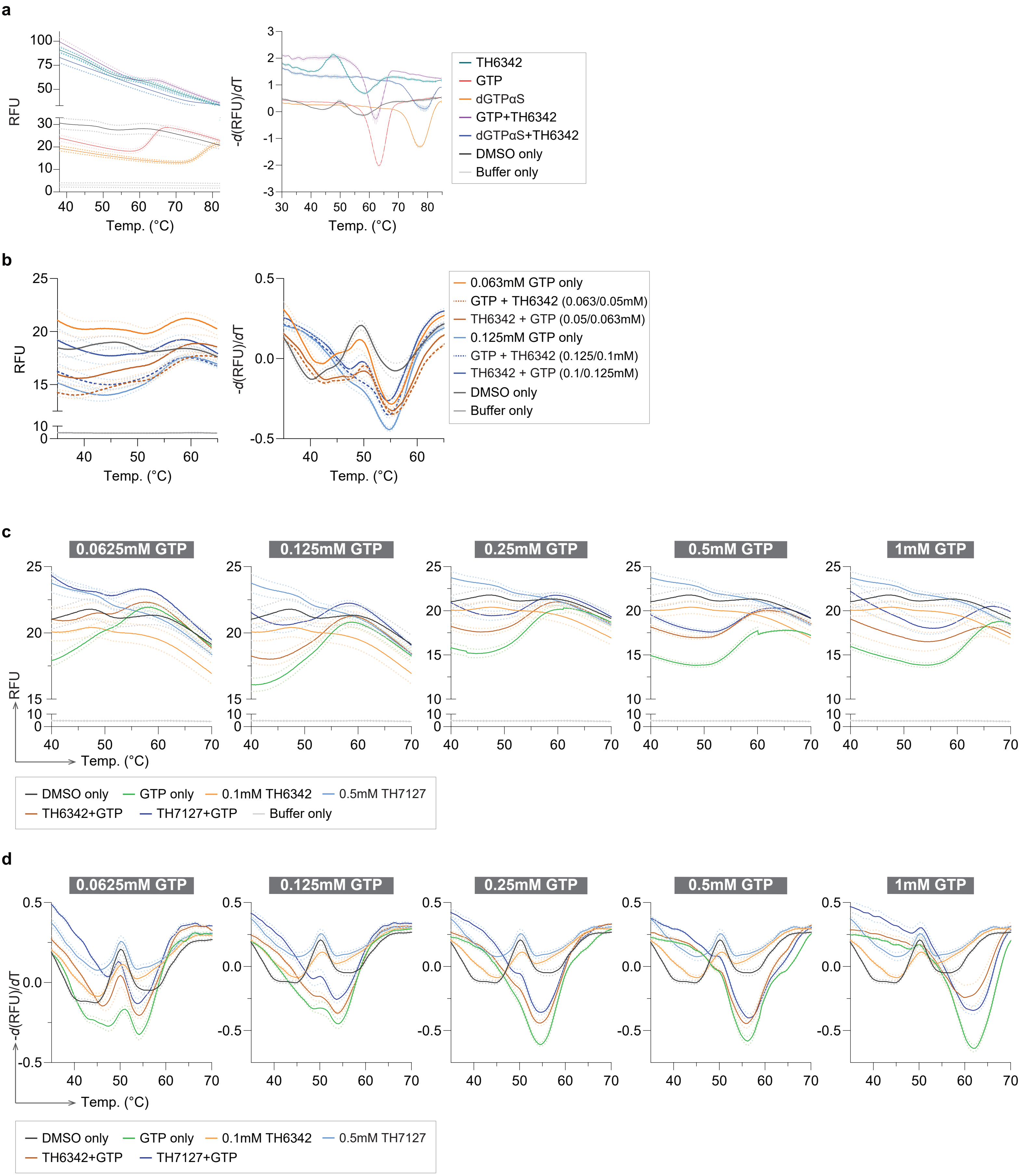
a
RFU
Temp. (°C)
-d(RFU)/dT
TH6342
GTP
dGTPαS
GTP+TH6342
dGTPαS+TH6342
DMSO only
Buffer only
b
0.063mM GTP only
GTP + TH6342 (0.063/0.05mM)
TH6342 + GTP (0.05/0.063mM)
0.125mM GTP only
GTP + TH6342 (0.125/0.1mM)
TH6342 + GTP (0.1/0.125mM)
DMSO only
Buffer only
c
0.0625mM GTP
0.125mM GTP
0.25mM GTP
0.5mM GTP
1mM GTP
DMSO only
GTP only
0.1mM TH6342
0.5mM TH7127
TH6342+GTP
TH7127+GTP
Buffer only
d
0.0625mM GTP
0.125mM GTP
0.25mM GTP
0.5mM GTP
1mM GTP
DMSO only
GTP only
0.1mM TH6342
0.5mM TH7127
TH6342+GTP
TH7127+GTP

**Supplementary Figure 6. Supplemental information of kinetic studies using the enzyme-coupled MG activity assay. a.** In the absence of putative inhibitors, SAMHD1 displayed minimal cooperativity in the enzyme-coupled MG activity assay with 25 μM dGTP as the self-activating substrate. Rate of reaction was fitted with either the Michaelis-Menten model (left panel) or an allosteric sigmoidal model ( right panel), using GraphPad Prism. Goodness of fit parameter $R^2$, as well as Hill coefficient ($H_n$) are shown. **b-c.** In the presence of TH6342 (b) or TH7127 (c), SAMHD1 displayed increasing levels of cooperativity in the enzyme-coupled MG activity assay with 25 μM dGTP as the self-activating substrate, supplementary to Fig. 4a and b, respectively. *Left panels*, Rate of reaction from Fig. 4a and b were re-fitted with the Michaelis-Menten model using GraphPad Prism. *Right panels*, Goodness of fit parameter $R^2$ from fitting with the Michaelis-Menten model were compared with those from fitting with an allosteric sigmoidal model as displayed in Fig. 4a and b. **d-e.** $K_{0.5}$ and $V_{max}$ values determined by fitting rate of reaction from Fig. 4a and b with an allosteric sigmoidal model (GraphPad Prism).

**Supp. Fig. 6**

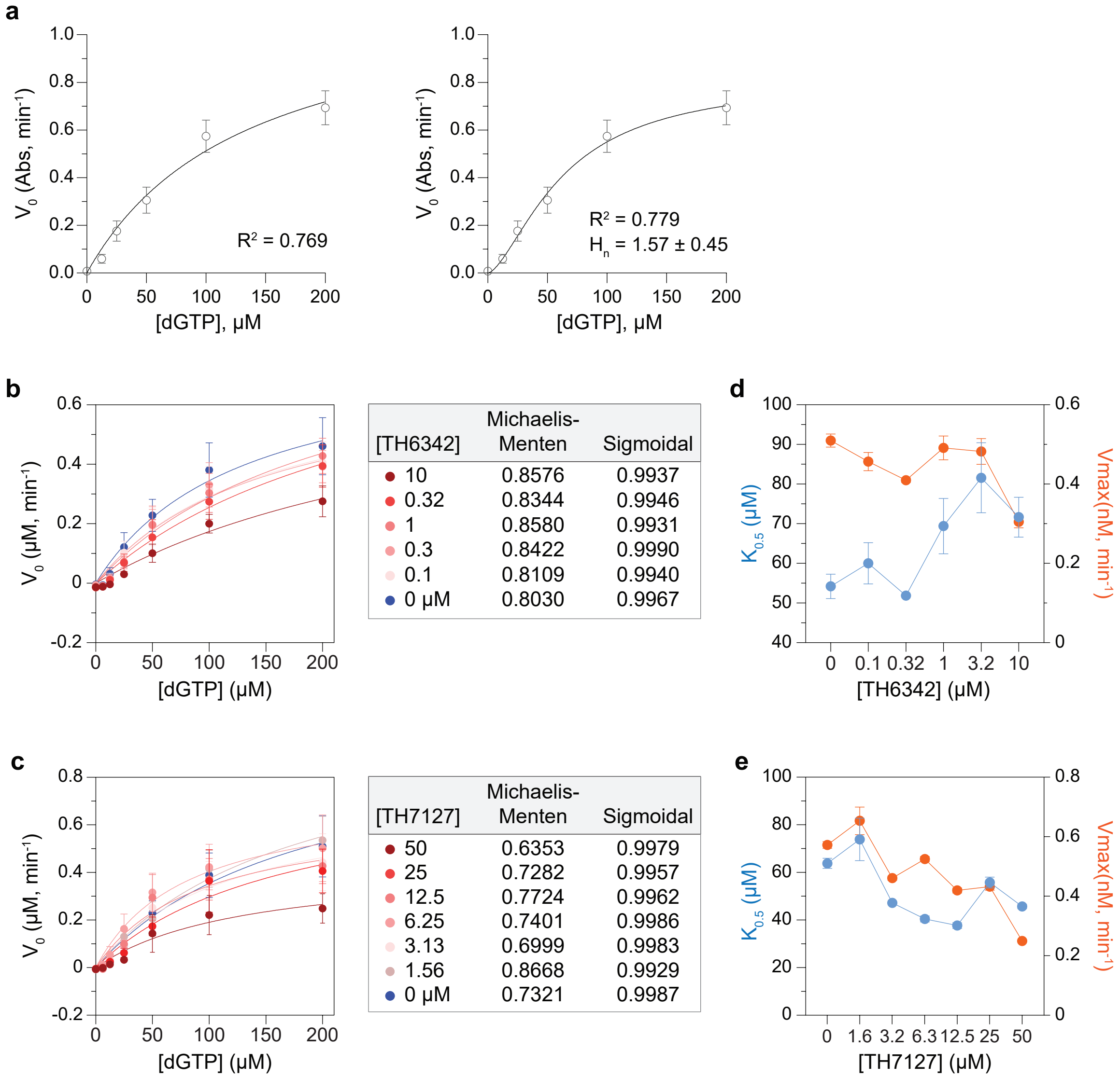


| [TH6342] | Michaelis-Menten | Sigmoidal |
|---|---|---|
| 10 | 0.8576 | 0.9937 |
| 0.32 | 0.8344 | 0.9946 |
| 1 | 0.8580 | 0.9931 |
| 0.3 | 0.8422 | 0.9990 |
| 0.1 | 0.8109 | 0.9940 |
| 0 µM | 0.8030 | 0.9967 |

| [TH7127] | Michaelis-Menten | Sigmoidal |
|---|---|---|
| 50 | 0.6353 | 0.9979 |
| 25 | 0.7282 | 0.9957 |
| 12.5 | 0.7724 | 0.9962 |
| 6.25 | 0.7401 | 0.9986 |
| 3.13 | 0.6999 | 0.9983 |
| 1.56 | 0.8668 | 0.9929 |
| 0 µM | 0.7321 | 0.9987 |

**Supplementary Figure 7. The b4NPP direct SAMHD1 enzymatic assay is linear under the specified conditions, as well as for the duration of experiments. a.** Establishment of linear range of detection of p-NP, product of b4NPP hydrolysis. Mean absorbance at 410 nm ($A_{410nm}$) ± SEM of n = 2 independent experiments each performed in triplicate are shown. *Left panel*, $A_{410nm}$ up to 5 mM, with the linear range of detection highlighted. *Right panel*, determined linear range of detection, where the mean $A_{410nm}$ values were fitted through a simple linear regression model (GraphPad Prism). $R^2$ value is shown. **b.** Titration of SAMHD1 concentrations. SAMHD1 of varying concentrations was incubated with 2 mM b4NPP, and $A_{410nm}$ was measured for up to 45 min. Mean $A_{410nm}$ ± SD of n = 2 independent experiments performed in triplicate or sextuplicate are shown. **c.** Titration of b4NPP concentrations. B4NPP at up to 4 mM was incubated with 0.5 μM recombinant SAMHD1 protein, and $A_{410nm}$ was measured for up to 45 min. Mean $A_{410nm}$ ± SEM of a representative experiments performed in triplicate are shown. *Left panel*, $A_{410nm}$ up to 45 min, with range of $A_{410nm}$ to determine $V_0$ highlighted. *Right panel*, $A_{410nm}$ up to 20 min, signals used to determine $V_0$ and subsequent kinetic study. **d.** Kinetic study of B4NPP hydrolysis by SAMHD1 under the specified conditions, in the presence of equivolume of DMSO (left panel) or TH7127 (right panel), supplementary to Fig. 4d. Mean $A_{410nm}$ ± SEM of n = 2 independent experiments performed in triplicate are shown, which were subsequently used to determine $V_0$ and mode of inhibition of TH7127.

**Supp. Fig. 7**

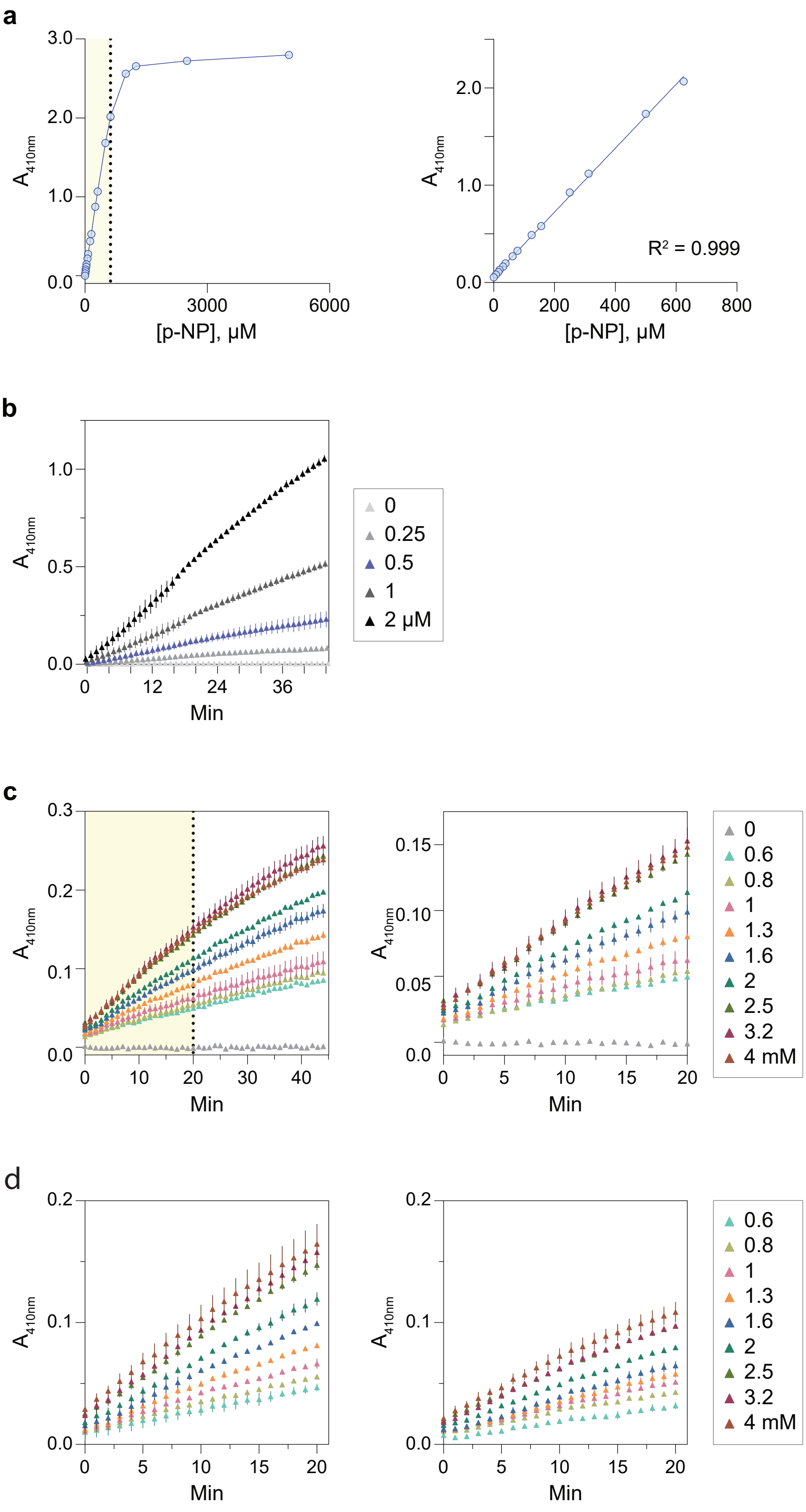

a
3.0
2.0
1.0
0.0
A410nm
0
3000
6000
[p-NP], μM
2.0
1.0
0.0
A410nm
R² = 0.999
0
200
400
600
800
[p-NP], μM
b
1.0
0.5
0.0
A410nm
0
12
24
36
Min
0
0.25
0.5
1
2 μM
c
0.3
0.2
0.1
0.0
A410nm
0
10
20
30
40
Min
0.15
0.10
0.05
0.0
A410nm
0
5
10
15
20
Min
0
0.6
0.8
1
1.3
1.6
2
2.5
3.2
4 mM
d
0.2
0.1
0.0
A410nm
0
5
10
15
20
Min
0.2
0.1
0.0
A410nm
0
5
10
15
20
Min
0.6
0.8
1
1.3
1.6
2
2.5
3.2
4 mM

**Supplementary Figure 8. Supplemental information of target engagement assays on cellular SAMHD1 protein. a.** Determination of screening temperatures in isothermal single-dose fingerprint CETSA experiments. Intact THP-1 cells or cell lysates were heated at increasing temperatures and were then analysed by Western blot for remaining soluble SAMHD1 protein. *Left panels*, representative Western blot images showing SAMHD1 and SOD-1 protein bands. *Right panel*, densitometry analysis of SAMHD1 signals, normalised to SOD-1 levels and then relative to samples heated at lowest temperatures. Mean relative SAMHD1 signals ± SEM of n = 2 independent experiments are shown. Melting curves are determined by curve-fitting mean relative SAMHD1 signals using a nonlinear regression model (Boltzmann sigmoidal, GraphPad Prism). Screening temperatures ($T_s$) are indicated. b. Optimisation of pronase concentration for SAMHD1 for DARTS experiments. Clarified THP-1 cell lysates were treated with increasing concentrations of pronase, followed by Western blot analysis to determine the optimal pronase concentration for DARTS study of SAMHD1.

Supp. Fig. 8

**a**

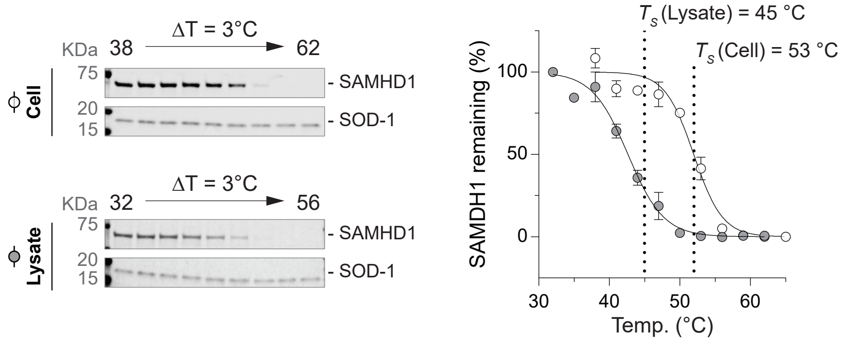
ΔT = 3°C
KDa 38 62
75
20
15
Cell
- SAMHD1
- SOD-1
ΔT = 3°C
KDa 32 56
75
20
15
Lysate
- SAMHD1
- SOD-1
T_s (Lysate) = 45 °C
T_s (Cell) = 53 °C
SAMDH1 remaining (%)
100
50
0
30
40
50
60
Temp. (°C)

**b**

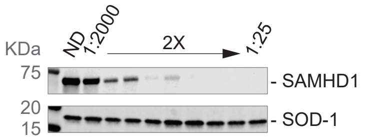
KDa
ND
1:2000
2X
1:25
75
20
15
- SAMHD1
- SOD-1

**Supplemental Figure 9.** The purity of recombinant SAMHD1 protein used in this study. Approximately 4 µg of SAMHD1 protein was analysed using SDS-PAGE, followed by Coomassie blue staining. Protein concentration was calculated from UV absorbance at 280 nm using a theoretical extinction coefficient.

**Supp. Fig. 9**

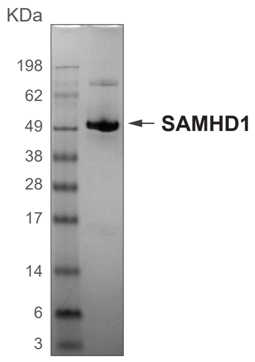

**/Source data Uncropped western blot images.**

**Source Data**

#### SAMHD1 melt curve

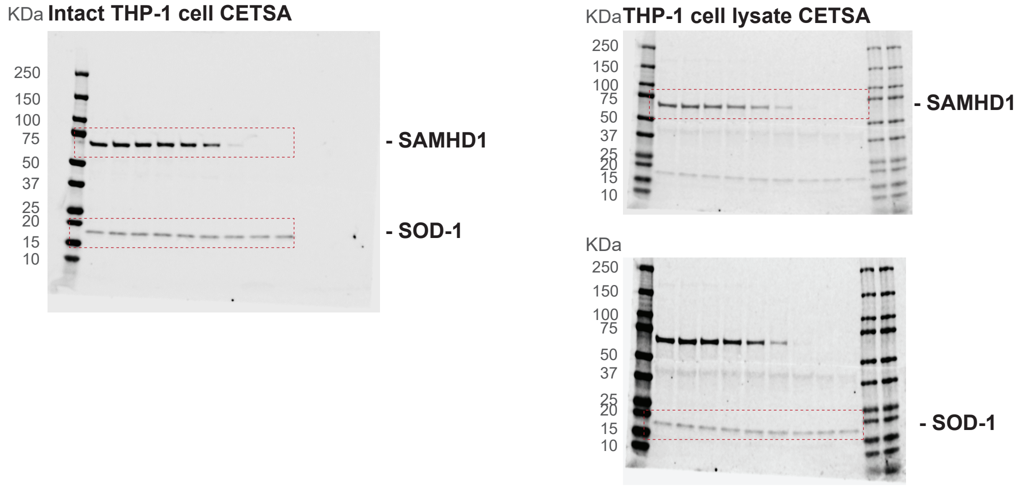


#### Intact CETSA with thymidine treatment

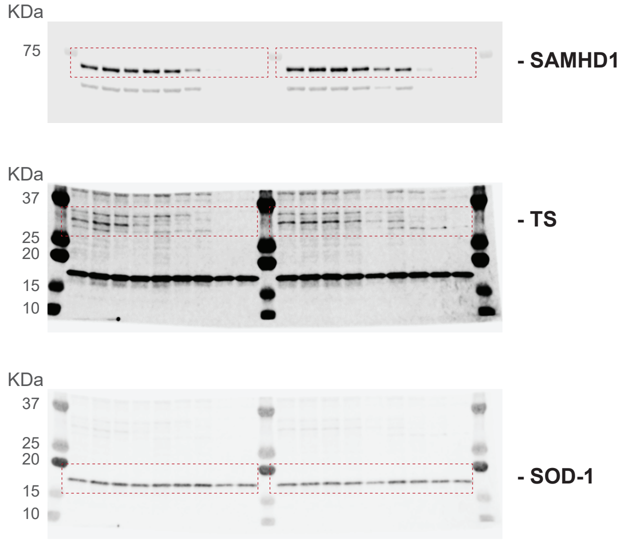


#### Lysate CETSA with dGTP treatment

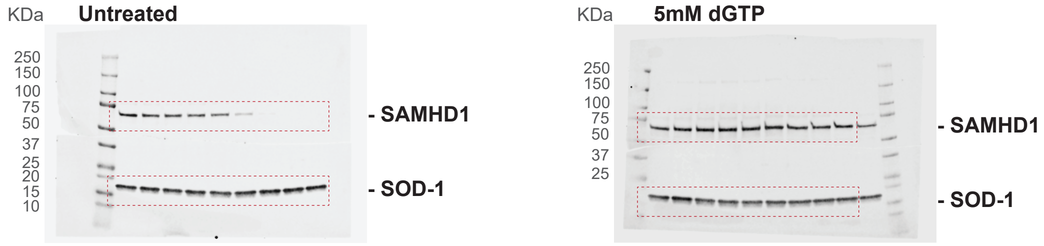


#### ITDRF SAMHD1 inhibitor

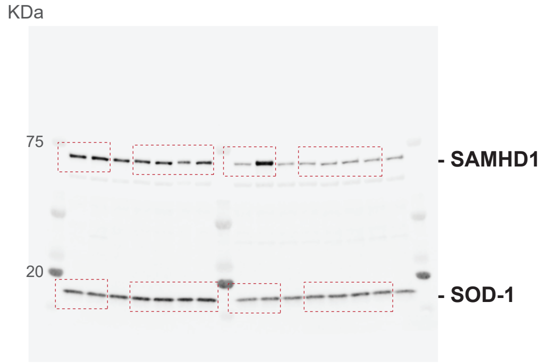

**Lysate CETSA with TH6342**

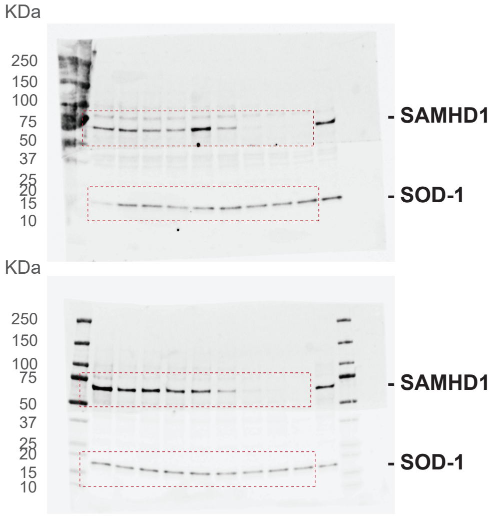


**SAMHD1 DARTS setup**

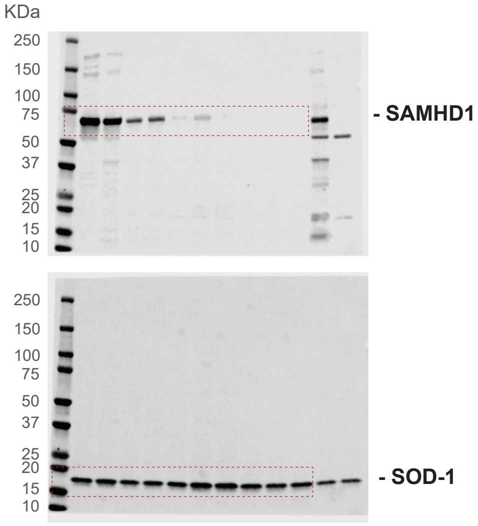


**SAMHD1 DARTS with dGTP**

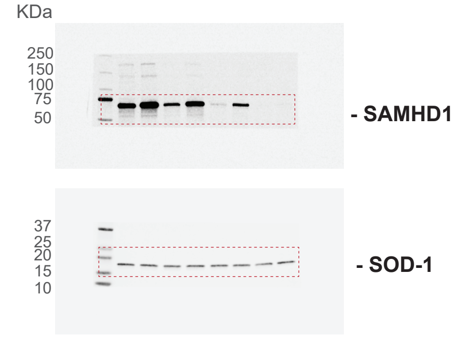


**SAMHD1 DARTS with TH6342**

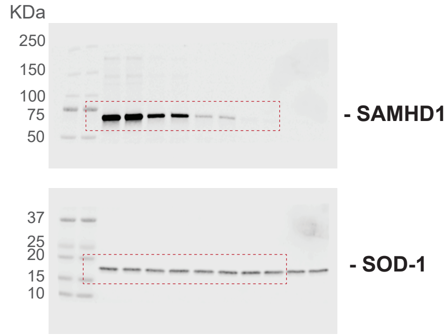


**SAMHD1 DARTS with TH7127**

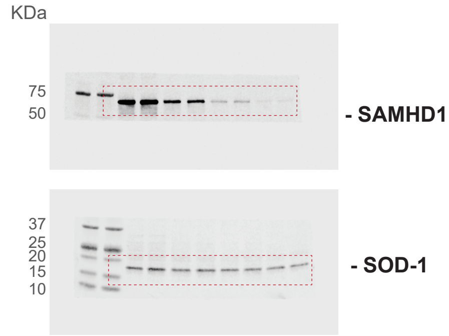

**Supplementary Table 1.** HTS screening campaign information

| Category | Parameter | Description |
|---|---|---|
| Assay | Type of assay | in vitro target based |
| | Target | Deoxynucleoside triphosphate triphosphohydrolase, SAMHD1 |
| | Primary measurement | Absorbance at 630 nm using a coupled enzymatic assay to detect inorganic phosphate (Pi) using the malachite green assay |
| | Key reagents | Human recombinant SAMHD1<br>The coupled enzyme inorganic pyrophosphatase was produced in *E. coli* by Protein Science facility at Karolinska institutet.<br>dGTP (Sigma-Aldrich D4010)<br>Stop reagent EDTA 7.9 mM<br>Malachite green detection reagents (Malachite Green, ammonium molybdate and Tween-20) |
| | Assay protocol | See supplementary information |
| | Additional comments | Protocol according to: Baykov, A. A., Evtushenko, O. A. & Avaeva, S. M. A malachite green procedure for orthophosphate determination and its use in alkaline phosphatase-based enzyme immunoassay. Anal. Biochem. 171, 266–270 (1988). Take care to ensure phosphate contamination in preparation of all reagents |
| Library | Library size | 17656 compounds |
| | Library composition | The library consists of a chemically diverse collection of compounds containing both commercial (Enamine, TimTec, Maybridge and ChemDiv) and internal compounds (donation from Biovitrum). The library includes a small fraction of compounds with known bioactivities, *e.g.* the Prestwick set, and a set of nucleosides from Barry Associates. |
| | Source | The screen was done based on plating of 10 mM DMSO solutions from Labcyte 384 LDV plates using an Echo 550 |
| | Additional comments | See below for further details on the composition of the Biovitrum derived compounds |
| Screen | Format | 384-well format<br>Assay plate: 384-well PS plate, Nunc 242757 |
| | Concentration(s) tested | Compound concentration at 5 µM, DMSO concentration at 0.1% |
| | Plate controls | Positive control: buffer only representing fully inhibited SAMHD1 enzyme (16 on each plate)<br>Negative control: uninhibited SAMHD1 enzyme (16 on each plate) |
| | Reagent/ compound dispensing system | Compound dispensing system: Echo 550 from Labcyte<br>Reagent dispensing system: FlexDrop IV from PerkinElmer<br>Multidrop from Thermo Scientific |
| | Detection instrument and software | Victor3 plate reader from PerkinElmer |
| | Assay validation/QC | Screen: Positive control: average absorbance 0.19, standard deviation 0.03. Negative control: average absorbance 0.52, standard deviation 0.01. Average Z´ factor/plate: 0.75.<br>QC also included monitoring of plate edge effects and distribution of the hits, with no corrections necessary |
| | Correction factors | Not applicable |

| | | |
|---|---|---|
| | Normalization | Data are normalized to the positive (100% inhibition) and negative controls (0% inhibition) on each plate and are expressed as % inhibition |
| | Additional comments | The screen was performed at Chemical Biology Consortium Sweden at Karolinska Institutet, Sweden |
| Post-HTS analysis | Hit criteria | Hit threshold = Average "% inhibition" of all test samples (0,36%) + 3 times standard deviation of all test samples (3*7.60%) = 23.15% |
| | Hit rate | 0.42% |
| | Additional assay(s) | Retesting of hits in 3 concentration hit confirmation experiment followed by a full concentration response experiments at 11 concentrations |
| | Confirmation of hit purity and structure | ID and purity analysis with LC-UV/MS detection |
| | Additional comments | |

**Supplementary Table 2.** Enzyme-coupled MG assay conditions.

| Enzyme | Coupled enzyme | Substrate | Enzyme conc. |
|---|---|---|---|
| MTH1 | PPase; 0.2U/ml | dGTP; 100 µM | 4.8 nM |
| NUDT15 | PPase; 0.2U/ml | dGTP; 100 µM | 8 nM |
| NUDT5 | BIP; 10U/ml | ADPR; 50 µM | 6 nM |
| NUDT12 | BIP; 10U/ml | β-NADH; 50 µM | 20 nM |
| NUDT22 | BIP; 10U/ml | UDP-glucose; 50 µM | 30 nM |
| ITPase | PPase; 0.45U/ml | ITP; 50 µM | 0.2 nM |
| dCTPase | PPase; 0.2U/ml | dCTP; 35 µM | 35 nM |
| dUTPase | PPase; 0.4U/ml | dUTP; 12.5 µM | 1.2 nM |

**Supplementary information on chemical synthesis**

All reagents and solvents were purchased from Sigma-Aldrich, Combi-Blocks, Thermo Fischer Scientific, or VWR and were used without purification. Unless otherwise stated, reactions were performed without care to exclude air or moisture. Analytical thin-layer chromatography was performed on silica gel 60 F-254 plates (E. Merck) and visualized under an UV lamp. Flash column chromatography was performed in a Biotage® SP4 MPLC system using Merck silica gel 60 Å (40–63 µm mesh). $^{1}H$ and $^{13}C$ NMR spectra were recorded on Bruker DRX-400 MHz and Bruker Avance 400 spectrometer. Chemical shifts are expressed in parts per million (ppm) and referenced to the residual solvent peak. For $^{1}H$ and $^{13}C$ measurements, the chemical shift is referred to an internal standard; the remaining protons or respectively the carbons of the corresponding deuterated solvent were used. Analytical LC–MS were performed on an Agilent MSD mass spectrometer connected to an Agilent 1100 system with: Method ST1090A3: Column ACE 3 C8 (50 × 3.0 mm); $H_2O$ (+ 0.1% TFA) and MeCN were used as mobile phases at a flow rate of 1 ml/min, with a gradient from 10% – 90% in 3 min; or Method B0597X3: Column Xterra MSC18 (50 × 3.0 mm); $H_2O$ (containing 10 mM $NH_4HCO_3$; pH = 10) and MeCN were used as mobile phases at a flow rate of 1 ml/min, with a gradient of 5% – 97% in 3 min. For LC–MS, detection was made by UV (254 or 214 nm) and MS (ESI+). Preparative LC was performed on a Gilson system using Waters C18 OBD 5 µm column (30 × 75 mm) with water buffer (a) 50 mM $NH_4HCO_3$ at pH 10 or b) 0.1% TFA) and acetonitrile as mobile phases using a flow rate of 45 ml/min. All final compounds were assessed to be >95% pure by LC–MS analysis.

***Precursor Synthesis: 6-bromo-N-(2-(pyridin-2-yl)ethyl)imidazo[1,2-a]pyrazin-8-amine***

250 mg (0.9 mmol) of 6,8-dibromoimidazo[1,2-a]pyrazine were dissolved in 5 mL Isopropanol and 1.1 eq. (119 µL, 0.99 mmol) 2-(pyridin-2-yl)ethanamine and 2 eq. (315 µL, 1.8 mmol) DIPEA were added. The reaction was stirred at 100°C overnight. The reaction mixture was evaporated and the crude was used for the synthesis of below compounds.

Alternatively, the precursor can be purified by silica column chromatography (10 g silica per 200 mg crude, ethyl acetate/isopropanol, gradient 0 to 5 %). The crude was dry-loaded onto silica in ethyl acetate/methanol (1:1). Target compound elutes at 3 % isopropanol in ethyl acetate. C13H12BrN5, M = 318.17 g/mol, LC-MS: $[M+H]^+$ 319; $^{1}H$-NMR (MeOD, 400 MHz): δ 8.34 (d, J = 4.67 Hz, 1H), 7.70 (s, 1H), 7.59 (dt, $J_1$ = 7.80 Hz, $J_2$ = 1.77 Hz, 1H), 7.57 (d, J = 1.01 Hz, 1H), 7.32 (d, J = 1.01 Hz, 1H), 7.21 (d, J = 7.80 Hz, 1H), 7.12 (td, $J_1$ = 6.32 Hz, $J_2$ = 0.90 Hz, 1H), 3.79 (t, J = 6.94 Hz, 2H), 3.04 (t, J = 6.94 Hz, 2H); $^{13}C$-NMR (MeOD, 100 MHz): 159.05, 148.31, 147.11, 137.19, 131.81, 131.85, 123.80, 122.48, 121.71, 115.39, 109.24, 40.21, 36.61; TLC (Ethyl acetate/isopropanol, 9:1): $R_f$ = 0.37.

***6-(2-chlorophenyl)-N-(2-(pyridin-2-yl)ethyl)imidazo[1,2-a]pyrazin-8-amine (TH6342)***

30 mg (0.094 mmol) of 6-bromo-N-(2-(pyridin-2-yl)ethyl)imidazo[1,2-a]pyrazin-8-amine was dissolved in 1 mL dioxane. Subsequently, 1.5 eq. (22.1 mg, 0.14 mmol) of (2-chlorophenyl)boronic acid, 0.2 eq. (21.8 mg, 0.018 mmol) tetrakis(triphenylphosphin)-palladium(0) and 126 µL of a 1 mM $Na_2CO_3$ solution were added and the reaction was refluxed

overnight. The reaction mixture was cooled to room temperature and filtered through Celite using MeOH. The solvents were evaporated and the crude product was purified over RP-HPLC to afford the desired product as a brown oil (3.3 mg, 9%). Alternative purification by silica column chromatography (Ethyl acetate/isopropanol, 10 g silica per 0.05 mmol substrate). The reaction crude was dry-loaded onto silica in acetone. Target compound elutes at 3 % isopropanol in ethyl acetate. C19H16ClN5, M = 349.104 g/mol; LCMS: $[M+H]^+$ 350; $^1$H-NMR (400 MHz, $CDCl_3$, δ = 7.27): 11.17 (brs, 2H), 9.04 (brs, 1H), 8.67 (dd, J = 5.7 Hz, 0.9 Hz, 1H), 8.13 (td, J = 7.8 Hz, 7.8 Hz, 1.6 Hz, 1H), 7.88 (s, 1H), 7.74-7.67 (m, 4H), 7.57-7.53 (m, 1H), 7.51-7.49 (m, 1H), 7.45-7.37 (m, 2H), 4.19 (brs, 2H), 3.50-3.47 (m, 2H) ppm.; $^{13}$C-NMR-DEPT (101 MHz, $CDCl_3$, δ = 77.2): 143.6, 142.6, 131.9, 130.3, 127.3, 126.8, 124.0, 109.7, 40.6, 33.6 ppm.

#### *6-(2-aminophenyl)-N-(2-(pyridin-2-yl)ethyl)imidazo[1,2-a]pyrazin-8-amine (TH7126)*

30 mg (0.094 mmol) of 6-bromo-N-(2-(pyridin-2-yl)ethyl)imidazo[1,2-a]pyrazin-8-amine was dissolved in 1 mL dioxane. Subsequently, 1.5 eq. (31.0 mg, 0.14 mmol) of 2-(4,4,5,5-tetramethyl-1,3,2-dioxaborolan-2-yl)aniline, 0.2 eq. (21.8 mg, 0.018 mmol) tetrakis-(triphenylphosphin)palladium(0) and 126 µL of a 1mM $Na_2CO_3$ solution were added and the reaction was refluxed overnight. The reaction mixture was cooled to room temperature and filtered through Celite using MeOH. The solvents were evaporated and the crude product was purified over RP-HPLC to afford the desired product as an orange solid (22.2 mg, 71%). C19H18N6, M = 330.1593 g/mol; LCMS: $[M+H]^+$ 331; $^1$H-NMR (400 MHz, $CDCl_3$, δ = 7.27): 8.59-8.57 (m, 1H), 7.62 (s, 1H), 7.59 (td, J = 7.7 Hz, 7.7 Hz, 1.8 Hz, 1H), 7.51 (d, J = 0.6 Hz, 2H), 7.33 (dd, J = 7.9 Hz, 1.1 Hz, 1H), 7.20-7.12 (m, 3H), 6.80-6.76 (m, 2H), 6.66 (t, J = 5.5 Hz, 1H), 4.58 (brs, 2H), 4.05 (q, J = 6.6 Hz, 2H), 3.19 (t, J = 6.6 Hz, 2H) ppm. ; $^{13}$C-NMR (101 MHz, $CDCl_3$, δ = 77.2): 159.1, 149.4, 147.3, 146.1, 140.1, 136.6, 132.1, 132.0, 129.4, 128.8, 123.4, 122.0, 121.6, 118.1, 116.9, 115.0, 108.0, 40.2, 37.4 ppm.

#### *6-(2-methoxyphenyl)-N-(2-(pyridin-2-yl)ethyl)imidazo[1,2-a]pyrazin-8-amine (TH7127)*

30 mg (0.094 mmol) of 6-bromo-N-(2-(pyridin-2-yl)ethyl)imidazo[1,2-a]pyrazin-8-amine was dissolved in 1 mL dioxane. Subsequently, 1.5 eq. (21.5 mg, 0.14 mmol) of (2-methoxyphenyl)boronic acid, 0.2 eq. (21.8 mg, 0.018 mmol) tetrakis-(triphenylphosphin)palladium(0) and 126 µL of a 1mM $Na_2CO_3$ solution were added and the reaction was refluxed overnight. The reaction mixture was cooled to room temperature and filtered through Celite using MeOH. The solvents were evaporated and the crude product was purified over RP-HPLC to afford the desired product as a green solid (11.7 mg, 34%). Alternative purification by silica column chromatography (Ethyl acetate/isopropanol, 10 g silica per 0.1 mmol substrate). Target compound elutes at 10 % isopropanol in ethyl acetate. C20H19N5O, M = 345.1590 g/mol; LCMS: $[M+H]^+$ 346; $^1$H-NMR (400 MHz, $CDCl_3$, δ = 7.27): 13.58 (brs, 2H), 8.98 (brs, 1H), 8.63 (d, J = 5.4 Hz), 8.42 (brs, 1H), 8.2 (td, J = 7.9 Hz, 7.9 Hz, 1.3 Hz, 1H), 8.05 (brs, 1H), 7.81 (d, J = 7.9 Hz), 7.73-7.71 (m, 2H), 7.60 (t, J = 6.5 Hz, 6.5 Hz, 1H), 7.41-7.37 (m, 1H), 7.09 (m, 1H), 7.00 (m, 1H), 4.23 (brs, 2H), 3.15 (s, 3H), 3.54-3.51 (m, 2H) ppm; $^{13}$C-NMR-DEPT (101 MHz, $CDCl_3$, δ = 77.2): 144.8, 141.9, 130.8, 130.7, 127.4, 124.4, 121.1, 111.3, 110.3, 55.6, 40.3, 33.2 ppm.

***N-(2-(pyridin-2-yl)ethyl)-6-(thiophen-2-yl)imidazo[1,2-a]pyrazin-8-amine (TH7528)***

27.5 mg (0.086 mmol) of 6-bromo-N-(2-(pyridin-2-yl)ethyl)imidazo[1,2-a]pyrazin-8-amine was dissolved in 1 mL dioxane. Subsequently, 1.5 eq. (16.6 mg, 0.13 mmol) of thiophen-2-yl boronic acid, 0.2 eq. (20.0 mg, 0.017 mmol) tetrakis(triphenylphosphin)palladium(0) and 115 µL of a 1mM $Na_2CO_3$ solution were added and the reaction was refluxed overnight. The reaction mixture was cooled to room temperature and filtered through Celite using MeOH. The solvents were evaporated and the crude product was purified over RP-HPLC to afford the desired product as a yellow oil (11.7 mg, 34%). C17H15N5S g/mol, M = 321.1048; LCMS: 322; $^{1}$H-NMR (400 MHz, $CDCl_3$, δ = 7.27): 8.68-8.66 (m, 1H), 7.83 (s, 1H), 7.81-7.79 (m, 1H), 7.54-7.53 (m, 2H), 7.48-7.45 (m, 2H), 7.35 (dd, J = 5.1 Hz, 1.1 Hz, 1H), 7.33-7.29 (m, 1H), 7.11-7.09 (m, 1H), 4.17-4.13 (m, 2H), 3.39 (t, J = 6.6 Hz, 2H) ppm; $^{13}$C-NMR-DEPT (101 MHz, $CDCl_3$, δ = 77.2): 146.9, 139.3, 127.9, 126.3, 124.9, 123.1, 122.4, 115.2, 40.5, 35.8 ppm.
